# Structural and Dynamic Insights Into α-Synuclein Dimer Conformations

**DOI:** 10.1101/795997

**Authors:** Joanna Zamel, Jiaxing Chen, Sofia Zaer, Paul David Harris, Paz Drori, Mario Lebendiker, Nir Kalisman, Nikolay V. Dokholyan, Eitan Lerner

## Abstract

Parkinson’s disease is associated with the aggregation of the protein α-synuclein. While α-synuclein can exist in multiple oligomeric states, the dimer has been a subject of extensive debates. Here, using an array of biophysical approaches, we demonstrate that α-synuclein *in vitro* exhibits primarily a monomer-dimer equilibrium in nanomolar concentrations and up to a few micromolars. We then use spatial information from hetero-isotopic cross-linking mass spectrometry experiments as restrains in discrete molecular dynamics simulations to obtain the ensemble structure of dimeric species. Out of eight structural sub-populations of dimers, we identify one that is compact, stable, abundant, and exhibits partially exposed β-sheet structures. This compact dimer is the only one where the hydroxyls of tyrosine 39 are in proximity that may promote dityrosine covalent linkage upon hydroxyl radicalization, which is implicated in α-synuclein amyloid fibrils. We propose that this α-synuclein dimer features etiological relevance to Parkinson’s disease.

## Introduction

α-Synuclein (αSyn) is a small soluble protein localized mainly in nuclei and in presynaptic terminals of dopaminergic neurons in the *substantia nigra pars compacta* (Recchia *et al*., 2004). In the brain, αSyn is involved in several stages of neurotransmitter trafficking, including the stabilization of dopamine vesicles and supporting fast and efficient formation of the soluble N-ethylmaleimide-sensitive factor attachment protein receptor (SNARE) complex (Burré *et al*., 2010). However, αSyn is involved in multiple disorders termed synucleinopathies, such as Parkinson’s disease (PD). One of the hallmarks of PD is the accumulation of αSyn amyloid-like fibrils in Lewy bodies (Recchia *et al*., 2004).

The protein is 140 amino acids long and can be divided into three peptide segments: 1) the N-terminal segment or domain (NTD; residues1-60), an amphipathic and lysine-rich region that plays critical roles in phospholipid attachment (Eliezer *et al*., 2001), 2) the non-amyloid-β component (NAC) segment (residues 61-95), a hydrophobic region that is involved in aggregation and fibrilization (Hashimoto *et al*., 2000; Uéda *et al*., 1993), and 3) the C-terminal segment or domain (CTD; residues 96-140), an acidic tail that contains multiple prolines and acidic amino acids that disrupt secondary structure formation (Ulmer *et al*., 2005).

αSyn attaches to different types of negatively charged membrane surfaces, including the ones that form dopamine vesicles in neurons (Bodner *et al*., 2010; Ferreon *et al*., 2009). Additionally, αSyn can spontaneously self-associate into oligomers. αSyn amyloid-like fibrils and certain oligomeric species are biochemical features of PD. Other oligomeric species have been documented, such as the α- helical stable tetramer (Bartels *et al*., 2011a; Binolfi *et al*., 2012; W. Wang *et al*., 2011). These different oligomeric species may support different biological functions.

In aqueous solutions, αSyn is intrinsically disordered. Therefore, αSyn exhibits features of misfolded proteins, such as high structural heterogeneity and conformational plasticity. Yet, oftentimes when αSyn binds to a biomolecular target, such as the outer leaflet of phospholipid membranes, or undergoes self-association, it tends to co-fold in a given pathway (Longhena *et al*., 2019).

αSyn aggregation kinetics include a long (tens of hours) delay, which is thought of as the time it takes for αSyn to form a fibrillization nucleus (Wördehoff & Hoyer, 2018). Since exposure of the hydrophobic residues of the NAC segment to the solvent is one factor that can help nucleate oligomerization in the fibril formation pathway, delay of the fibrillization nucleation can be attained via αSyn forms that sterically occlude the NAC from exposure to the solvent. In the free-form monomer, the CTD and NTD sterically interact to shield the NAC from being exposed to the solvent, and by that protect the NAC segment from aggregation-prone interactions with the NAC segment of other αSyn molecules (Levitan *et al*., 2011). However, there may be other αSyn forms, including small oligomers, which may act to sterically occlude the NAC.

αSyn self-associates and some of its self-associated species lead to pathophysiological-promoting conditions, such as amyloid-like fibrils or toxic oligomers. Therefore, it is of utmost importance to study the mechanism by which αSyn self-associates, and specifically the earliest stages of oligomerization. Logically, the earliest stage of self-association would be the result of dimerization. It is noteworthy that αSyn dimers and trimers have previously been implicated with neurotoxicity (Coelho-Cerqueira *et al*., 2013; Outeiro *et al*., 2008). Additionally, the αSyn dimer would serve as the smallest and most elementary self-associated species. The monomer-dimer interaction could be considered controversial, as several studies had demonstrated that αSyn exists as a monomer in solution (Fauvet, Mbefo, *et al*., 2012; Theillet *et al*., 2016; Weinreb *et al*., 1996), or as a tetramer (Bartels *et al*., 2011b).

Previously, there have been computational and experimental works providing evidence for the existence of αSyn dimers using ion mobility mass spectrometry (MS), paramagnetic resonance enhancement (PRE) nuclear magnetic resonance (NMR), spin-label NMR as well as potential models of the αSyn dimer from molecular dynamics (MD) simulations (Janowska *et al*., 2015; Kang *et al*., 2012; Lan-Mark & Miller, 2022; T. Zhang *et al*., 2017). Nevertheless, structure models of the αSyn dimer that are based on experimental and computational works integratively combined have not yet been introduced.

Therefore, studying the structural features of an αSyn dimer is important for understanding how self-association starts in αSyn at the first place. Additionally, it is hypothesized that specific inter-molecular interactions within αSyn oligomers can act as seeds of aggregation, especially irreversible interactions, such as covalent bonds between pairs of amino acid side chains of two αSyn subunits (Pivato *et al*., 2012). One such covalent linkage that is related to the stabilization of the fibrillization pathway is dityrosine covalent linkages (Pivato *et al*., 2012; Souza *et al*., 2000; Takahashi *et al*., 2002; van Maarschalkerweerd *et al*., 2015). Dityrosine linkages can form only after hydroxyl radicalization of tyrosines, which can occur in specific stress conditions, implicated in PD (Gaeta & Hider, 2005). Therefore, upon elucidation of structure models of the αSyn dimer, inter-molecular proximal tyrosines may serve as a signature of the potential for this covalent linkage to occur under such stress conditions.

Several experimental and computational studies have previously focused on the αSyn dimer (Illes-Toth *et al*., 2015; Kayed *et al*., 2020; Li *et al*., 2019; Lv *et al*., 2015; Mane & Stepanova, 2016; Pivato *et al*., 2012; Tsigelny *et al*., 2007; Y. Zhang et al., 2018). Using high-speed atomic force imaging, Zhang *et al*. (Y. Zhang et al., 2018) have revealed the αSyn dimer exists in several different configurations, at low spatial resolution: (i) with one compact globular subunit and another expanded and dynamic, (ii) with both subunits being compact and globular, and (iii) with both subunits being expanded and flexible.

Here, we perform an array of experiments that show fresh αSyn molecules are predominantly monomers and dimers *in vitro* at concentrations of few micromolars (µM) at most, which is below the known concentrations at which αSyn exhibit efficient aggregation that leads to fibril formation (Wördehoff & Hoyer, 2018). Using single-molecule photo-isomerization related fluorescence enhancement (smPIFE; commonly known as protein-induced fluorescence enhancement) (Chen, Zaer *et al*., 2021; Hwang *et al*., 2011; Hwang & Myong, 2014a; Zaer & Lerner, 2021), we find that dimer formation is accompanied by local structural changes in the vicinity of specific residues, at the subunit level, pointing towards local structural differences between the monomer and dimer αSyn.

Encouraged by the results, we performed hybrid structural modeling of the αSyn dimer, using hetero-isotopic cross-linking MS (CL-MS) (Gaber *et al*., 2019; Lima *et al*., 2018; Walker *et al*., 2014) as experimentally-derived restraints for discrete MD (DMD) simulations, to elucidate structure models of the αSyn dimer. We report the features of the structure models of different conformations of the αSyn dimer.

## Results

### MALS: αSyn monomer-dimer mixture

We perform analytical anion exchange (AIEX) chromatography coupled to multi-angle light scattering (MALS), yielding a direct readout of the molecular mass of the different species in solution with high detection sensitivity and minimal chemical modifications or perturbations (Amartely *et al*., 2018). AIEX-MALS of *wt*-αSyn yields a single elution peak with an average molecular mass of 21.6±0.8 KDa, which is larger than that of a monomer (14.4 KDa) and smaller than that of a dimer (28.8 KDa; Fig. 1A). *wt-*αSyn eluted as a single peak using the monoQ 1 mL column (Fig. S1). Based on the MALS results, we propose that the elution peak contains a mixture of primarily αSyn monomers and dimers, leading to the resolved average molecular mass between a monomer and a dimer. Assuming the αSyn dimer is unstable, the mixture of a monomer and a dimer can be a manifestation of an existing monomer-dimer equilibrium. However, these results are based on AIEX performed against salt gradient, and hence could be suspected to be an outcome of the presence of elevated salt concentrations. Therefore, to strengthen our findings we further perform validation of the monomer-dimer mixture suggestion using size exclusion chromatography (SEC)-MALS (Figs. 1B, S2 and S3). Importantly, previous SEC-MALS studies of αSyn had shown that using the same SEC resin and similar procedure as ours, αSyn trimers and tetramers may exist, but elute earlier than a monomer-dimer major elution peak (Burré *et al*., 2013). Our results show a chromatogram with similar results, just without evidence for the separate trimer-tetramer elution peak: after a first elution peak of low amounts of early-formed aggregates, we record one main elution peak with an average molecular mass (identified directly from MALS) larger than a monomer and smaller than a dimer (Fig. 1B), with values almost identical to the values resolved in AIEX-MALS. We can rule out the presence of trimers and tetramers in our mixture by relying on the work of Burre’ *et al*. Therefore, the ensemble of αSyn can be described as a mixture of primarily monomers and dimers.

**Fig. 1:**
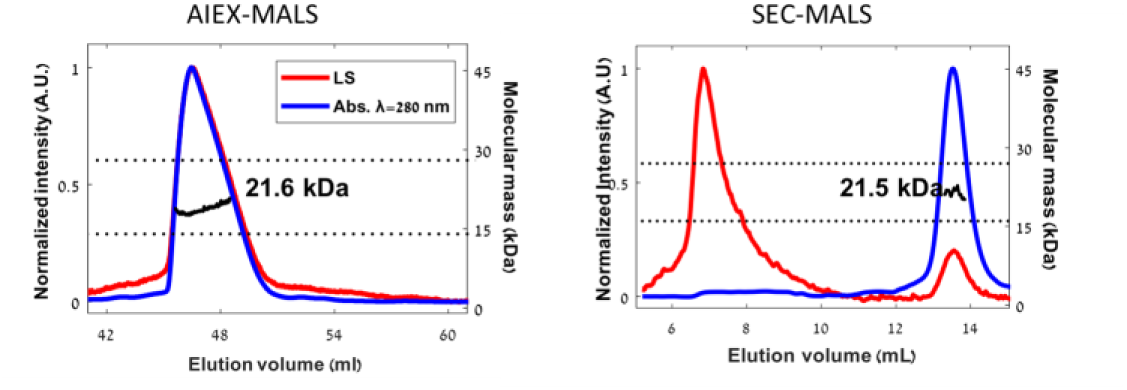
MALS of *wt*-αSyn elutes as a single peak with a molecular mass larger than αSyn monomer. **A**. AIEX-MALS, the absorption at 280 nm (blue) and the light scattering intensity (red) were used for the calculation of MALS- derived molecular weight of 21.6±0.8 KDa. The elutant had a molecular mass value between that expected for an αSyn monomer and that expected for a dimer (dotted grey lines). **B.** SEC-MALS using a Superdex 200 column was performed after injection of the *wt*-αSyn, and both the absorption at λ=280 nm (blue) and the light scattering intensity (red) were monitored. The early elution peak was characteristic of aggregates exhibiting high light scattering intensity and low protein absorption. The late elution peak was characteristic of a species larger than a monomer, due to the MALS-derived molecular mass of 21.4±5 KDa that is larger than the monomer molecular mass of 14.4 KDa.

### Western Blot of cross-linked αSyn: monomers and dimers mixture

After characterizing the presence of αSyn dimers at µM concentrations, our goal would be to perform cross-linking mass spectrometry (CL-MS) with the goal of studying their structural organization. However, before doing so, we would like to check that the cross-linking does not lead to higher-order oligomers. Therefore, we conducted cross-linking experiment using bis-sulfosuccinimidyl suberate (BS^3^) cross- linking reagent in two different concentrations of *wt*-αSyn, followed by western blot (WB) with αSyn antibody and SDS-PAGE (see *methods*).

The WB (Fig. 2) shows the presence of a band at ∼14 KDa and another weaker band of ∼28 KDa in both concentrations indicating the presence of monomers and dimers of αSyn, respectively. At 10 µM of αSyn the dimer band is stronger and more noticeable, while in at 5 µM the band is fainter, but present (also see Fig. S4). As the performed WB is only qualitative and not a quantitative method, we have no indicator of the dimer fraction. Two important findings are shown: (i) as the concentration of αSyn increases in the few µM range, the fraction of the αSyn dimer increases, and (ii) the WB together with the AIEX/SEC-MALS rule-out the presence of higher order αSyn oligomers. These results allow us, later in this work, to perform CL-MS analyses of inter- and intra-molecular crosslinking in the αSyn dimer.

**Fig. 2:**
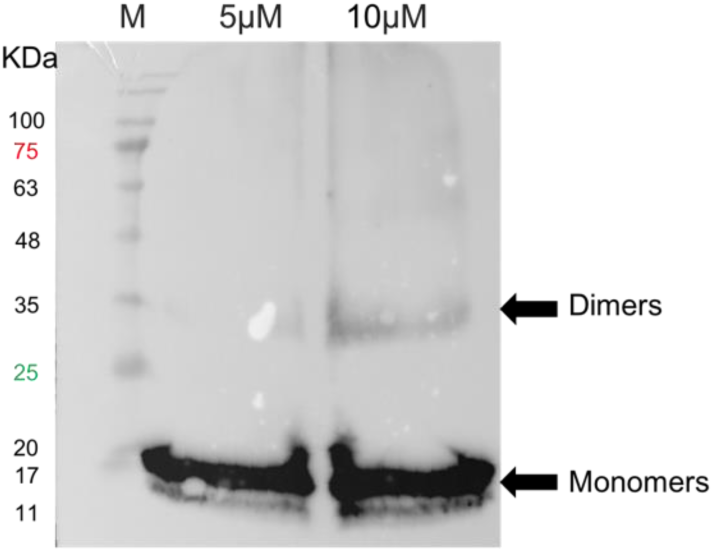
Western blot of cross-linked *wt*- αSyn. The left column is marker (M), middle and right columns are αSyn in two concentrations, 5 and 10 µM after cross- linking using 1 mM BS^3^. In both αSyn concentrations two bands are present at ∼14 KDa and 28KDa. Note that the dimer band in the 5 µM column is hard to observe. For a clearer result, see Fig. S4 for the same result after 40 seconds exposure time.

### Fluorescence-based assays: small-scale self-association equilibrium

For assessing αSyn dimerization as a self-association process, we perform several inter-molecular Förster resonance energy transfer (FRET) and single-molecule photo-isomerization-related fluorescence enhancement (smPIFE) measurements (Figs. S5-S9).

Although steady-state inter-molecular FRET measurements clearly indicate inter- molecular FRET occurring between 100 nM αSyn labeled at residue 39 with a donor dye (ATTO 488) and 100 nM αSyn labeled at residue 39 with an acceptor dye (ATTO 647N) (Fig. S5), we strive to test these results at different conditions including lower αSyn concentrations, for which a standard steady-state fluorimeter is not sensitive enough. To do so, we record acceptor fluorescence decays after donor excitation. If a fraction of the donor- and acceptor-labeled αSyn molecules under measurement is within a dimer and exhibiting FRET, a clear signature should appear in the donor-excited acceptor fluorescence decays as a delayed fluorescence decay, which should be enhanced as a function of the dimer fraction, hence as a function of increasing αSyn concentration.

We tested two pairs of donor- and acceptor-labeled proteins: (i) a mixture of αSyn labeled with either ATTO 488 (donor) or ATTO 647N (acceptor) at the thiol introduced by cysteine at position 39, an experiment we shortly term 39-39, and (ii) a mixture of αSyn labeled with the donor dye ATTO 488 thiol introduced by cysteine at position 39 and by the acceptor dye ATTO 643 at the thiol introduced by cysteine at position 140, an experiment we shortly term 39-140.

Fig. S6 summarizes the results of the 39-39 time-resolved inter-molecular FRET experiment. The donor-excited acceptor normalized fluorescence decays pertain, even at low αSyn concentrations. This is due to a small yet nonnegligible fraction of acceptor fluorescence following direct excitation of the acceptor attained with the excitation source used for donor excitation, which is low (Fig. S7), as well as a very low contribution from donor fluorescence leakage into the acceptor detection channel. Nevertheless, the number of molecules exhibiting the delayed fluorescence decay, hence the fraction of molecules exhibiting inter-molecular FRET, is low at low αSyn concentrations. Overall, the higher αSyn concentration is, the more delayed the donor-excited acceptor fluorescence decays are (Fig. S6). We have also performed fluorescence correlation spectroscopy (FCS)-FRET measurements of 39-39 to show the effect of inter-molecular FRET at low αSyn concentrations (see Fig. S8 and SI for additional details).

While these were the results for the 39-39 time-resolved intermolecular FRET experiment, for 39-140 the results show a similar pattern however at a different αSyn concentration range (Fig. 3). Above a certain αSyn concentration (300 nM), no change is observed to the decay rates, indicating a saturation level is reached. Below 300 nM, it is noticeable that the lower αSyn concentration is the faster decaying are the donor-excited acceptor fluorescence decays, whilst no effect is observed in the acceptor-excited acceptor fluorescence decays.

**Fig. 3:**
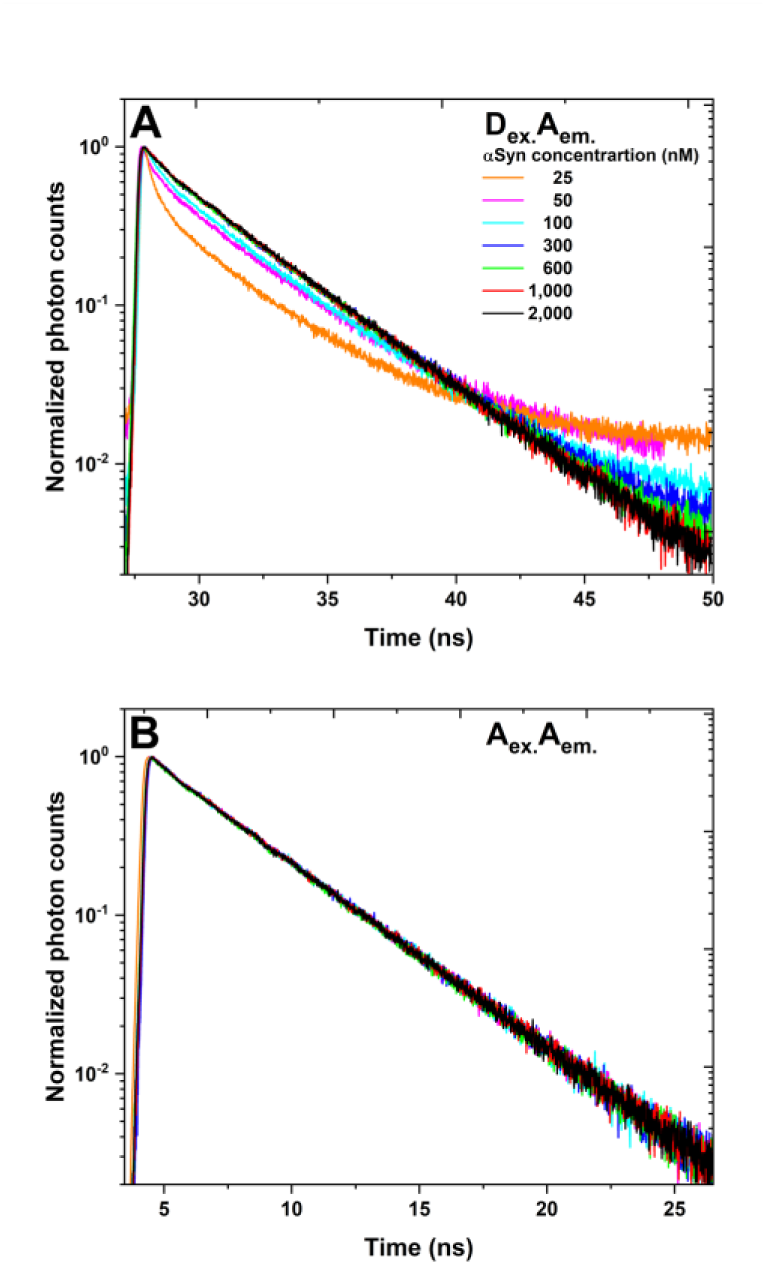
A signature of intermolecular FRET in acceptor fluorescence decays, decreases with decreasing αSyn concentration (of a mixture of αSyn labeled at either residue 39 or residue 140). **A**. The acceptor fluorescence decays, following donor excitation, were measured in donor- and acceptor-labeled αSyn mixtures at different αSyn concentration. The fluorescence decay curves exhibit at least two components: a fast component, that is most probably due to leakage of the donor fluorescence red edge into the acceptor detection channel (see Fig. S7) and a slow component, indicative of acceptor excitations indirectly that were delayed in the donor excited state prior to the FRET event. The amplitude of the slow component is decreasing as the concentration of αSyn decreases. **B**. As a control, the acceptor fluorescence decays, following acceptor excitation, which seems to show no change in the decay rates as a function of αSyn concentration.

Our steady-state, time-resolved, and FCS-FRET (Figs. 3, S5, S6, S8) measurements of inter-molecular FRET together with the IEX/SEC-MALS and WB results presented earlier, demonstrate a monomer-dimer equilibrium exists. Therefore, results of smPIFE measurements (Fig. S9) are expected to be related to the monomer-dimer changes (see results in SI). In summary, the smPIFE results report on the change in site-specific steric obstruction levels as the concentration of *wt*-αSyn increases, pointing towards structural rearrangements the αSyn is undergoing while forming the dimer. To ensure that the *wt*-αSyn titration results in dimer formation and not higher-order self-associated species, we correlated the sCy3 mean fluorescence lifetimes with their burst durations (Fig. S10). Burst durations are indicative of the diffusion through the confocal spot, and hence are proportional to the hydrodynamic radii of the measured molecules, which is a connection that has been shown rigorously (Hagai & Lerner, 2019). Indeed, our measurements report on changes in sub-populations of sCy3 fluorescence lifetimes that are correlated with a 30-50% increase in the burst durations in each dye-labeled residue, which agrees with the previously reported increase in the hydrodynamic radius of monomeric αSyn versus dimeric αSyn (Li *et al*., 2019). Therefore, we expect the αSyn dimer to be represented by multiple distinct conformations.

### Hetero-isotopic CL-MS: retrieving multiple spatial restraints to support hybrid structural modelling

To the best of our knowledge, a high-resolution structure model of the αSyn dimer is not available. The main reason that hinders solving the structure by NMR or X-ray crystallography is the oligomeric and structural heterogeneity of αSyn. Therefore, there is a need to use a different approach to overcome these difficulties. Here, we introduce cross-linking mass-spectrometry (CL-MS) followed by discrete molecular dynamics (DMD) as an integrative structural biology approach to model the structure of the αSyn dimer.

CL-MS has emerged as a robust method for studying protein structures and assemblies (Gonzalez-Lozano *et al*., 2022; Graziadei & Rappsilber, 2022; Kolesnikova *et al*., 2018; Koukos & Bonvin, 2020; Merkley *et al*., 2014; Na & Paek, 2020; Yang & Tang, 2019). Recently, CL-MS has been used to study small disordered proteins, such as the free-form αSyn monomer, and assisted in elucidating its ensemble structure (Chen, Zaer *et al*., 2021). One of the limitations of CL-MS is the inability to distinguish between inter- and intra-molecular cross-links in homodimers. One way to circumvent this limitation is by metabolic labeling of one subunit with the heavy isotope, ^15^N, while the other monomer remains as the natural light ^14^N isotope (Gaber *et al*., 2019; Lima *et al*., 2018; Walker *et al*., 2014). The mass shift of the heavy isotope allows the distinction between inter- or intra-molecular cross-links (Fig. 4). In this approach, we mix the ‘heavy’ and the ‘light’ monomers and introduce the cross-linking reagent. Besides the heavy-heavy and light-light peptides that may originate from inter- or intra-molecular interactions, we will obtain heavy-light cross-linked peptides that can only originate from inter-molecular interactions (Fig. 4). Additionally, light-light or heavy-heavy cross-linked residues, which have not appeared in the list of light-heavy cross-linked residues, are suggestive of intra- molecular cross-links. Nevertheless, based on MALS measurements the homodimer is proposed to be in equilibrium with the dissociated monomer. Therefore, it is hard to tell which intra-molecular cross-link comes from the dimer subunit and which comes from the dissociated monomer. Therefore, it is important to work in conditions in which the homodimer is enriched, and the amount of dissociated monomer is minimal (see *Methods*).

**Fig. 4:**
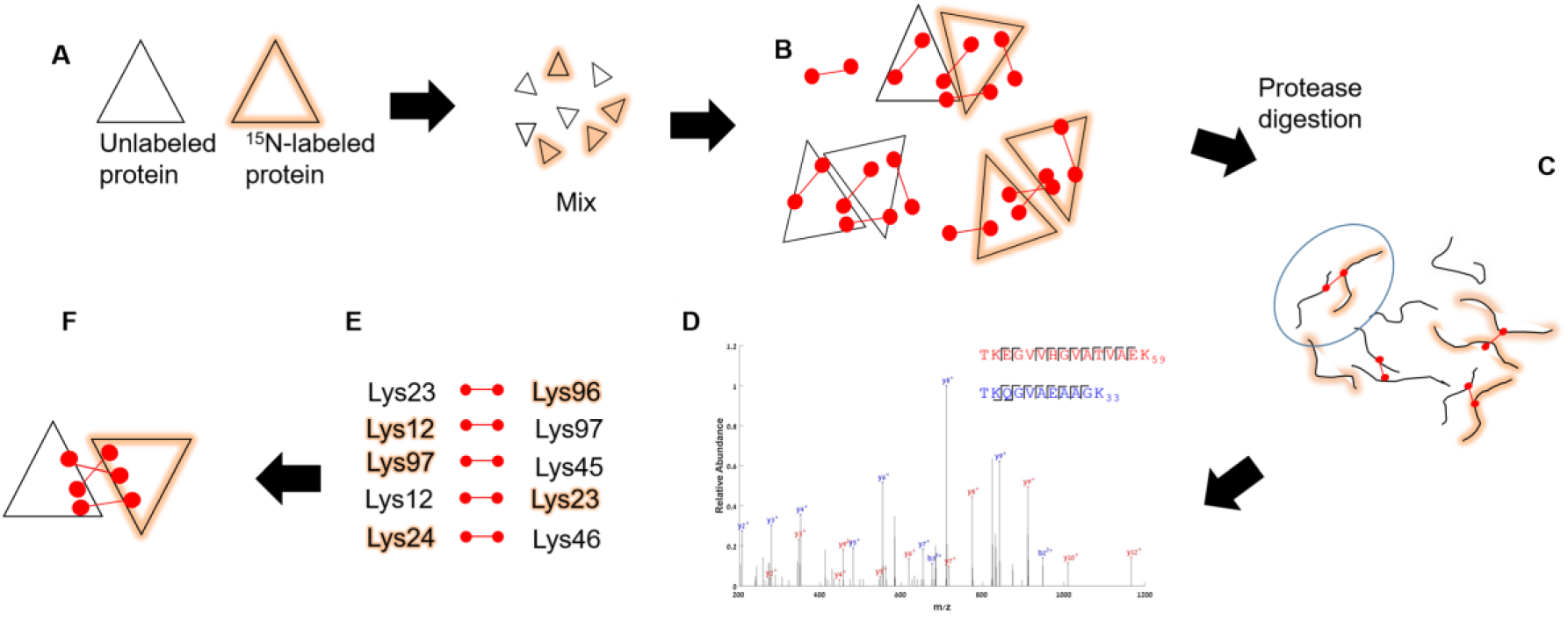
Hetero-isotopic CL-MS workflow. The ^14^N-^15^N CL-MS workflow. **(A)** ^14^N- and ^15^N-labeled α-syn are mixed. **(B)** Incubation with cross-linkers (red dumbbells). Covalent inter and intra molecular cross-links in the homodimer (within the cross-linker length) **(C)** Complexes are digested via a protease, ending with peptides linked with the cross-linker. Pairs of hetero- isotopic peptides can originate only from inter-molecular cross-links. Similarly, cross-links between a pair of homo-isotopic peptides that are not identified as inter-molecular, are intra-molecular **(D)** The digested peptides analyzed by mass-spectrometry and the spectra help identify pairs of cross-linked residues. Differently cross-linked pairs of peptides with different nitrogen isotope contents yield different distinguishable peaks. **(E)** Hundreds of cross-links can be identified from a large complex. **(F)** The cross-links are converted into distance restraints/constraints, which drive computational modeling.

We conduct three different experiments on an equimolar mixture of either ^14^N-^14^N, ^15^N -^15^N, or ^14^N -^15^N α-Syn with a total concentration of 2.5 µM. For the experiment we used two bifunctional cross-linkers: i) the homo-bifunctional amine-reactive linker BS^3^ with <30Å Cα-Cα distance, and ii) the hetero-bifunctional amine-carboxyl reactive coupler 4-(4,6-dimethoxy-1,3,5-triazin-2-yl)-4-methyl-morpholinium chloride (DMTMM), a zero-length linker <16Å Cα-Cα distance (Leitner *et al*., 2014). We prepared the sample by standard preparation protocols (Slavin & Kalisman, 2018), digested the proteins with trypsin or GluC, which cleaves after lysine or after glutamate and aspartate, respectively. We then perform the mass spectrometry analyses (see *Methods*).

Using BS^3^, we identify a total of 121 cross-links, of which 103 are homo-isotopic and 18 are hetero-isotopic (see *supplementary spreadsheet CL-MS*). The estimated false discovery rate (FDR) is 5% (see methods). The majority of BS^3^ inter-molecular cross-links are between the NTD and the NAC segments, due to the lack of lysines in the CTD (Fig. 5A). To gain better cross-link coverage of the C-terminal we apply DMTMM and obtain 23 inter-molecular cross-links (Fig. 5B).

**Fig. 5:**
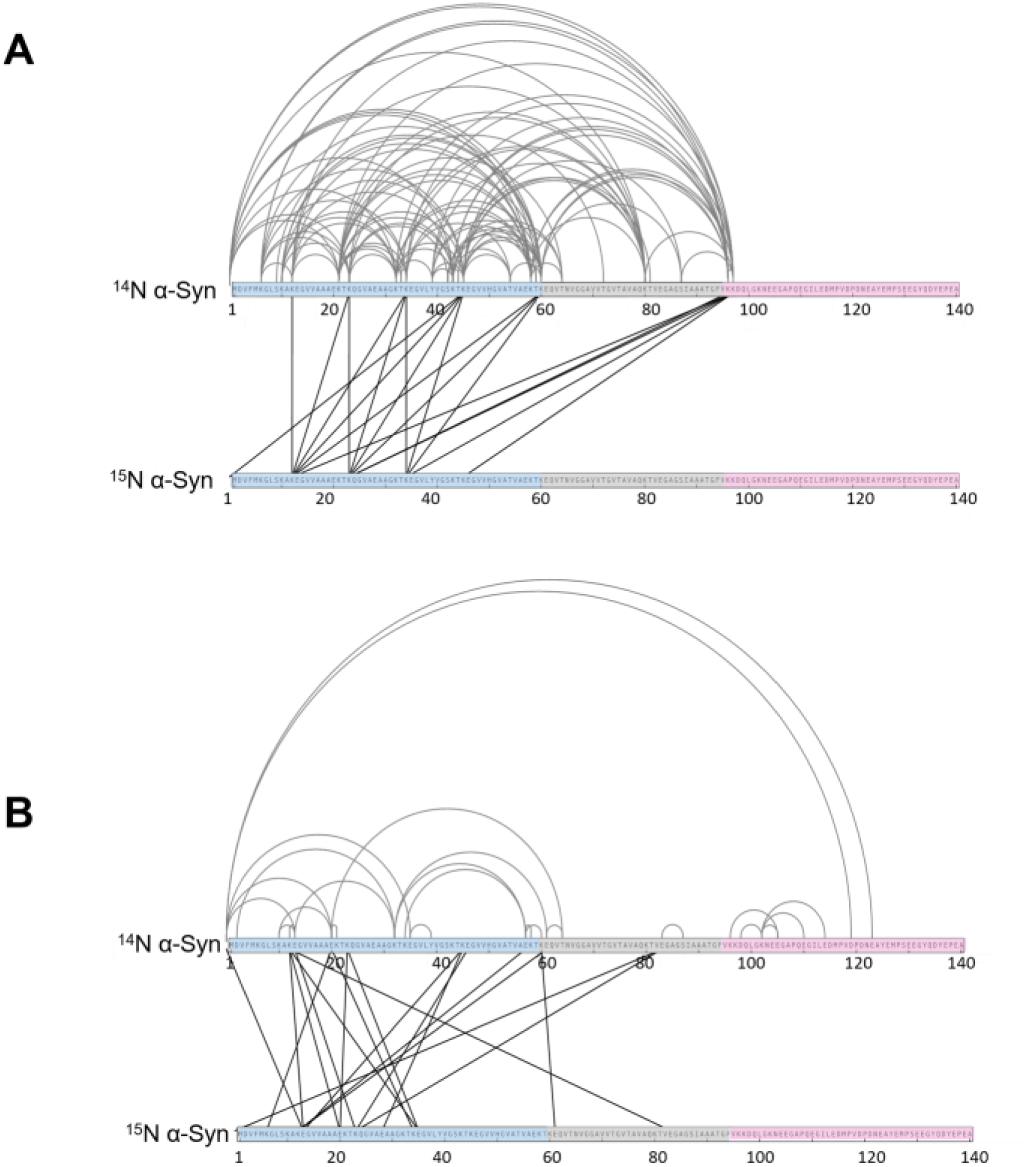
Hetero-isotopic CLMS of αSyn dimer. Grey – intra-molecular cross-links; Black – inter-molecular cross- links. (**A)** BS^3^ with Trypsin digestion. (**B**) DMTMM with either Trypsin or GluC digestion

To gain short range spatial information, potentially regarding proximities to the CTD segment, we use DMTMM and digest using GluC. However, we could not identify any inter-molecular cross-link with the CTD segment (Fig. 5B). This indicates that there are no short-range proximities between one subunit and the CTD segment of the other subunit in the dimer. In contrast, we identify intra- molecular cross-links between the CTD segment and the other segments within the subunit.

Remarkably, the pattern of the inter-molecular interactions between the NAC and NTD segments within the dimers captured by cross-linking include interactions between the NTD segment of one subunit and the NAC segment of the other subunit. These results indicate a specific fold is expected between the NTD and NAC segments in the dimer.

In addition to understanding the residue-residue proximities in the homodimer, gained from CL-MS information, we are able to acquire multiple non-redundant inter-molecular spatial restraints. The CL-MS attained spatial restraints are sufficient to direct the experimentally-restrained DMD simulations to form dimers in the simulation and to result in structure models, which exhibit partial agreement with the experimental data, and ones not biased by the cross-linking distance ranges (see restraints designed for DMD simulations in Fig. S11).

### Experimentally-restrained DMD simulations: structure models of αSyn dimer conformations

We elucidate αSyn dimer structure models by cross-linking restraint-guided DMD simulations (for details, see *Methods*). To analyze the dimer structures, we choose the most-populated conformations and perform agglomerative hierarchical clustering (see *Methods*), which divides the structural ensemble into eight clusters of different compactness, as judged by radii of gyration (Rg), and different energies (Fig. 6A). Structures in each cluster are highly similar as the pairwise root mean square deviation (RMSD) in each cluster is <9 Å (Table S2). The RMSD between clusters (Table S3) is >14 Å, indicating that structures among different clusters are distinct from each other. Then, we extract the dimer structure models and validate them against the cross-linking data (Fig. 6B). Specifically, we calculate the fraction of structures within each cluster, which satisfy the inter-molecular cross-linking restraints (Table S4; cluster-based distance distribution histograms deposited over Zenodo (Lerner *et al*., 2021a)). If a majority of structures in a cluster exhibit the inter-residue distances values within the Cα-Cα distance range the cross-linkers cover (BS^3^ with Cα-Cα distance <30 Å; DMTMM with Cα-Cα distance <16 Å) (Leitner *et al*., 2014), these cross-links are confirmed as agreeing with the structure model of the cluster. All 121 intra-molecular and 36 inter-molecular cross-links are tested against our dimer structure models (almost all intra-molecular cross-links agree well with the structural data; we focus on the inter-molecular cross-linking data). Following this, clusters 1, 2, and 4 contain more structures, which are validated by inter-molecular cross-links, than the other clusters (Fig. 6B). That is because we design the restraints to energetically favor distances covered by the cross-linkers, but not to abolish potential longer distances, and hence not to bias the simulations.

**Fig. 6:**
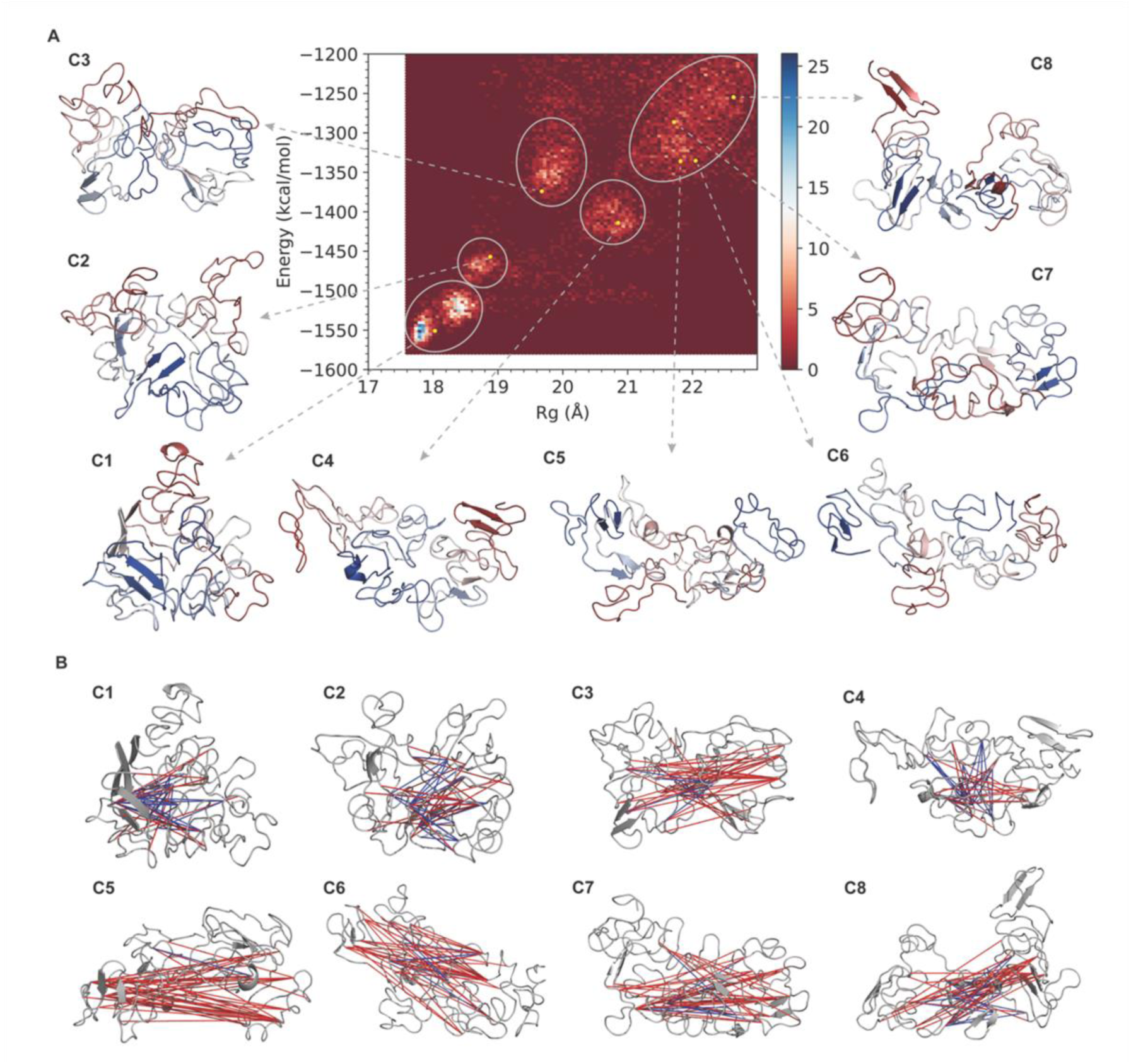
The eight structural sub-populations or clusters of the αSyn dimer predicted by cross-linking- restrained DMD simulations. (A) The hexbin plot of energy versus radius of gyration (Rg) for simulated dimer structures and the centroid structure of each cluster (C1-C8). The structures are marked from blue (NTD segment) to red (CTD segment). (B) The centroid structures of the eight clusters. Each line in the structure indicates the inter-molecular cross- links used for DMD simulations. Blue line means that over 50% of the structures within the corresponding cluster are validated by this constraint, while red line implicates that this constraint is satisfied by less than 50% of the structures for each cluster. The exact values of the levels of agreement of structure models with cross-linking data can be found in table S4.

The results of our smPIFE experiments identify several sub-populations of the αSyn dimer (Fig. S9). To investigate if the simulation-predicted dimer structures include different sub-populations or not, we calculate the solvent accessible surface area (SASA) of the accessible volumes of Cy3 when it is attached to residues 26, 39, 56, and 140 within each monomer for all the eight clusters. The higher the SASA value is, the more accessible is the residue. Importantly, clusters with no histograms point to residues that are fully buried and inaccessible. Residue 26 has three sub-populations in both αSyn subunits (Fig. S12). Residue 39 contains three sub-populations with different degrees of accessibility (SASA values centered at ∼1,000, 1,500, and 2,500) in chain A and B (Fig. S13). Residue 56 exhibits two SASA sub-populations in one subunit and three in the other subunit (Fig. S14). Residue 140 exhibits three SASA sub-populations in either one of the subunits (Fig. S15). Overall, the different sub-populations represented by our simulated dimer structures are in accordance with the smPIFE data.

Two tyrosine residues could form a dityrosine covalent linkage, if they are spatially close enough and if their hydroxyl groups are radicalized. This bond formation stabilizes homo-oligomeric αSyn complexes. Moreover, a Y39-Y39 dityrosine linkage has previously been reported as occurring in αSyn and stabilizing its later oligomeric species, some of which in the aggregation *on-pathway*, potentially acting as stable seeds of aggregation (Souza *et al*., 2000; van Maarschalkerweerd *et al*., 2015). Another dityrosine covalent linkage has been reported for αSyn in Y125-Y125 (Takahashi *et al*., 2002). To find out if αSyn dimers have pairs of tyrosines from the two subunits that are within interaction distances to form this covalent bond, we calculate the distance distributions between hydroxyl groups of all permutations of pairs of such tyrosine groups (e.g., involving tyrosine residues 39, 125, 133 and 136) in each of the clusters. Out of these permutations, the only pair of tyrosines, which exhibits the short distances required for interaction, is the Y39-Y39 pair, and only in cluster 1 (Fig. S17). The distances between hydroxyl groups of tyrosine residues in cluster 1 are significantly smaller than those in the other clusters, with >50% of its structures exhibiting distances <6 Å. These dimer structures have the potential to form a covalent bond between tyrosine residues, if at least one of their hydroxyl groups have undergone radicalization, which can covalently stabilize the dimer structure. Therefore, cluster 1 is of particular interest. In addition, cluster 1 has the lowest energy compared to the other clusters (Fig. 6A) and is the most compact (Fig. 6A), the most populated cluster and it is the cluster that exhibits the highest level of agreement with the CL-MS data. The NTD segment as well as the NAC and CTD segments in one subunit of the centroid of cluster 1 form β-sheet structures (Fig. 7), with a partial degree of surface exposure in both NTD and NAC segments (Fig. S16). Therefore, out of all structure models of the αSyn, the one described by cluster 1 is of most interest.

**Fig. 7:**
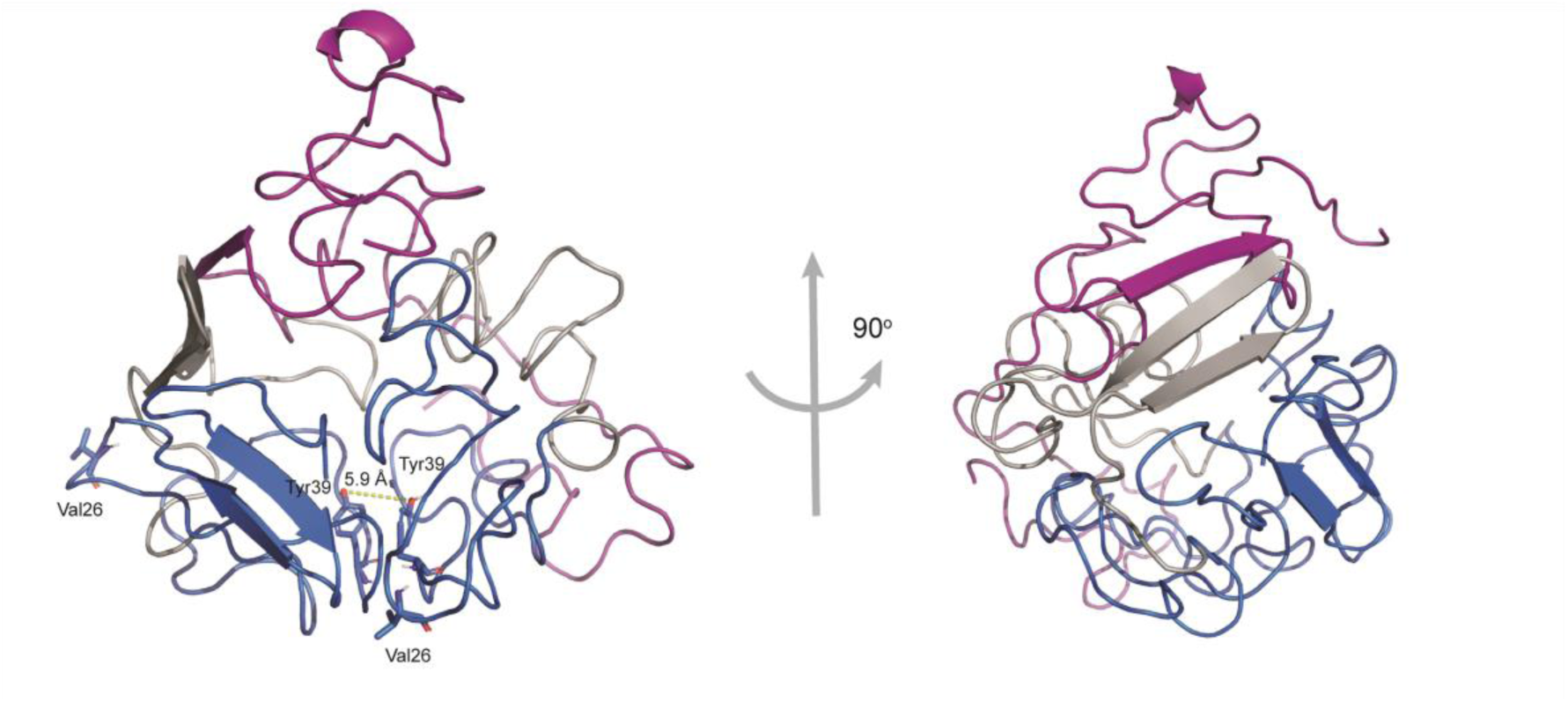
The cartoon representation of the centroid structure of cluster 1. The NTD, NAC, and CTD segments are highlighted in blue, grey, and magenta, respectively. Note that while the NTD and NAC segments of both subunits in close proximity, the CTD segments of both subunits are far from each other, in line with the hetero-isotopic CL-MS results in Fig. 5. Tyrosine 39 of both subunits are shown, with the short proximity between their hydroxyl groups. Additionally, it is clear that Tyrosine 39 is not accessible and buried, and is well characterized by the high mean nanotime sub-population for this residue in Fig.S9. Residue 26, one of the other hand is exposed and accessible in both subunit, and hence is well characterized by the low mean nanotime sub-population for this residue in Fig.S9.

## Discussion

We performed AIEX-MALS and SEC-MALS, WB, inter-molecular FRET assays and smPIFE measurements, to demonstrate a monomer-dimer mixture, and potentially a monomer-dimer equilibrium exists in αSyn at the concentration range of a few μM. Therefore, when probing a solution of freshly-thawed recombinant αSyn with a nominal concentration of a few μM, a significant number of dimers are formed (Figs. 1,2). To gain insight into dimer structures, we perform hetero-isotopic CL-MS, which retrieves inter- and intra-molecular spatial information on the dimer and use the cross-linking data as restraints in DMD simulations (Dokholyan, 2020).

Our experimentally-restrained DMD simulations retrieve an ensemble of structure models clustered in eight structural sub-populations or clusters (C1-8). It is noteworthy that since the RMSD values within clusters C1-8, perhaps except for C4, are lower than 6 Å (Table S2), it is relatively safe to infer structural features from the inspection of the centroid structures of the retrieved clusters. The energies of the centroid structures of these clusters increase (Fig. 6A) as the number of amino acids in the dimerization interface decreases (Table S5). Additionally, this trend appears after analyzing the non-covalent interactions at the dimer interface, using the PPCheck (Sukhwal & Sowdhamini, 2013) server (see Table S5 and *supplementary spreadsheet PPCheck*). Among all the clusters, the centroid structure model of C1 has several important features being the most populated cluster in the simulation (17.3%) with the lowest energy (Fig. 6A), the lowest stabilizing energy (Table S5), and it appears to have the highest number of residues in the interface between the subunits (155 residues; Table S5). Furthermore, the centroid structure of C1 is the most compact (Fig. 6A). Overall, it seems that among all eight clusters, C1 is the most stable αSyn dimer species, with multiple inter-molecular interactions (Table S5). Interestingly, the structure of one subunit of the dimer in C1 exhibits partially buried NTD and NAC segments and a more solvent-exposed CTD segment. Additionally, the structure of the other subunit in the C1 dimer exhibits NTD and NAC segments that are even more buried than their counterparts in the first subunit, and again a relatively solvent-exposed CTD segment. This observation indicates that either C1 is not on the NAC-induced aggregation pathway, which requires solvent exposure of the NAC segment, or that C1 is an intermediate, where later changes in conformation, upon further oligomerization or covalent stabilization may expose the NAC. Additionally, β- sheet structures are found in this subunit of C1, in its NTD segment and in its NAC and CTD segments. These β-sheet structures of the stable and abundant dimer conformation C1 exhibit partial accessibilities, and therefore might serve in aggregation pathways alternative to the one promoted by the solvent-exposed NAC segment β-sheet structure. Recently, Lan-Mark & Miller have introduced potential αSyn homo-dimer structure models based on MD simulations (Lan-Mark & Miller, 2022). In agreement with their conclusion, various polymorphic αSyn dimers might be obtained using different MD simulation methods. Herein, we introduce structures that are based not only on theoretical modeling, but also on solution-based experiments and using them as restraints for structural modeling.

Importantly, we do not make any assumption as to the initial structures in the DMD simulations other than starting from fully extended conformations, and we start collecting information from the simulations only after we prove it has equilibrated.

Covalent stabilization of adjacent αSyn subunits within aggregates has been previously reported to enhance the formation of stable aggregates and seeds of aggregation (Pivato *et al*., 2012; van Maarschalkerweerd *et al*., 2015). Specifically, the formation of Y39-Y39 dityrosine linkages between two αSyn subunits in an aggregate was characterized in that respect (Mukherjee *et al*., 2017) and a structural model was proposed to explain the observed results (Souza *et al*., 2000). However, other dityrosine covalent linkages, such as Y125-Y125 were also implicated (Takahashi *et al*., 2002). We examine the inter-molecular distances between the hydroxyl groups of all tyrosine pairs against the structures of each cluster out of the eight identified clusters. Only one cluster, in only one tyrosine pair, Y39-Y39, exhibits distances within interaction distances (Fig. S17), which upon hydroxyl radicalization can generate such dityrosine linkages. Such tyrosine hydroxyl radicalization can occur as a result of oxidative or nitrative stress (Souza *et al*., 2000). All the characteristics of C1 raise the question of whether the αSyn dimer could be stabilized by covalent dityrosine linkages, and if it can, will it stay in the C1 conformation, or could it undergo conformational changes after dityrosine linkages covalently stabilize the dimer in such a way that the NAC will later be fully exposed to the solvent to promote aggregation.

C2 is low in the energy, however not as low as C1 (Fig. 6A), and its centroid structure includes fewer residues in the dimerization interface than C1 (Table 5). Additionally, the centroid structure of C2 is less compact than that of C1 (Fig. 6A). Like in C1, the centroid structure of C2 exhibits β-sheet structures, some of which include multiple long β strands (Table S6). C3 and C4 are higher in energy relative to C1 and C2 (Fig. 6A), their centroid structures include fewer residues within the dimerization interface (Table S5), and they are more expanded relative to the centroid structures of C1 and C2 (Fig. 6A). However, the centroid structure of C4 is expected to be more stable than that of C3, with lower total stabilizing energy values (Table S5). None of the structure models of C2-4 exhibit Y39-Y39 proximity to accommodate the potential formation of a dityrosine covalent linkage.

The existence of multiple αSyn dimer conformations is warranted by the existence of multiple mean nanotime sub-populations in time-resolved smPIFE experiments (Fig. S9), signifying the existence of dimers with different local structures in the vicinity of the labeling sites (Hwang & Myong, 2014b). Moreover, these different local-structure related sub-populations undergo structural changes at times slower than single-molecule burst durations, hence at milliseconds or slower. Nevertheless, it is noteworthy that at the moment we cannot exclude the possibility that each mean nanotime sub-population is the result of time-averaging of transitions faster than typical burst duration timescales, better known as *within-burst dynamics*. We have recently developed an analysis approach such dynamics in single-molecule FRET experiments (Harris *et al*., 2021), and will soon extend this approach to analyze time-resolved smPIFE within-burst dynamics.

In a different work that used intermolecular smFRET to remarkably track single αSyn oligomers of different types, αSyn oligomers of different types and sizes were identified (Cremades *et al*., 2012). In our work, we report predominantly monomers and dimers of αSyn. Is there a discrepancy? The excellent work of Cremades *et al*., used a stock of high concentration αSyn that was shaken and incubated and then at each time point (hours), diluted to single-molecule concentrations for data acquisition. In such conditions, it was expected that with time stable oligomers and aggregates will form. The results shown in this work were performed as fast as possible on freshly-thawed aliquots, and hence the conditions used in our work are not expected to induce αSyn aggregates efficiently. Additionally, at concentrations of dye-labeled αSyn as low as 3 nM, FCS-FRET reports only minimal FRET (Fig. S8). Even so, for the sake of comparison, since we can perform smFRET measurements, we tested intermolecular smFRET of donor- and acceptor-labeled αSyn directly diluted to 100 pM concentrations from freshly-thawed αSyn stocks. The results included only negligible fractions of bursts with inter-molecular FRET (Fig. S18). In summary, the two works are not comparable due to the different conditions used, where in our work we were able to gain insights predominantly about monomers and dimers of αSyn, as our data indicate.

Back to the integrative structural modeling of the αSyn dimer, although the restraints derived from hereto-isotopic CL-MS play a pivotal role in the experimentally-restrained DMD simulations of the αSyn dimer, one major limitation is the inability to report whether the intra-molecular cross-links originate from an αSyn monomer or the subunit of the dimer, since both are found in equilibrium. In that respect, it is noteworthy that the concentrations of αSyn used in hetero- isotopic CL-MS are ones, in which the fraction of the dimer is high, as can be seen from our chromatography coupled to MALS, WB and inter-molecular FRET titration measurements. Additionally, the DMD restraints were designed to give less energetic weight for the intra-molecular cross-links and more for the inter- molecular cross-links (Fig. S11, compare panels A and B).

The structure models of the αSyn dimer contribute to the understanding of the oligomerization pathway of αSyn, which has been extensively studied, both for recombinant αSyn and for αSyn purified from natural sources (Andreasen *et al*., 2015; Cremades *et al*., 2012; de Franceschi *et al*., 2011; Du *et al*., 2020; Ingelsson, 2016; Li *et al*., 2019; Salveson *et al*., 2016). Nevertheless, to date, the mechanism is not fully understood. To enhance our understanding of the first step in αSyn oligomerization, we studied the smallest known αSyn oligomer, which marks the first inter-molecular self-association encounter - the dimer. αSyn dimers have been previously studied using SDS-PAGE, atomic force microscopy (AFM) (Zhang *et al*., 2018), and reporting on the formation of the dityrosine covalent linkage in induced dimers (Takahashi *et al*., 2002; van Maarschalkerweerd *et al*., 2015). Using PRE-NMR, in line with our findings showed variety of conformations of homo-dimers. It has also been shown by ion-mobility spectrometry combined with electrospray ionization MS-MS (ESI-IMS-MS) that αSyn in physiological conditions exists as monomer and dimer with the dimer population estimated at only ∼5%- 10% (Janowska *et al*., 2015).

Importantly, in our work we heavily rely on the mass estimation from SEC- and AIEX-MALS and the values that are higher than the molecular mass of a monomer. Could it be that instead of observing a mixture of monomers and dimers as we suggest, the mixture also includes fractions of trimers, tetramers or larger oligomers? Luckily, it has been shown that in SEC-MALS trimers and tetramers of aSyn, which we do not recover, elute earlier than the monomer-dimer elution peak, using the same SEC column and purification procedure as the one we used (Burré *et al*., 2013). Therefore, our results together with these studies strengthen our estimate of an exclusive monomer-dimer mixture.

Our structure models describe αSyn dimers without modifications, although in the cytosol αSyn is reported to be constitutively N-terminus acetylated (Burré *et al*., 2013). A number of studies have indicated that N-terminus acetylation of αSyn either has no effect or has only minor effects on fibril structural properties (Bartels *et al*., 2014; Fauvet, Fares, *et al*., 2012) and effects on the fibrillization kinetics and aggregation propensities (Bartels *et al*., 2014; Watson & Lee, 2019), while other reports indicated contradicting results (Fauvet, Fares, *et al*., 2012; Kang *et al*., 2013). Therefore, the effects of N-terminus acetylation on αSyn fibrillization are under debate. As for dimers of αSyn, we did not identify studies explaining the importance of N-terminus acetylation on αSyn dimers.

For now, it is acceptable that small forms of αSyn in the cell are mainly found as a disordered monomer (Fauvet, Mbefo, *et al*., 2012). Our findings may open the possibility for the presence of αSyn dimers as other small forms of αSyn in the cell as well, however neither we nor anyone else have provided evidence for that.

## Summary

Altogether, we identify structural models of the dimer, some of which are interesting from the functional point of view, whether they are or are not in the NAC-inducing aggregation pathway. These structural models will be the input for a follow-up work, in which we intend to design point mutations that will selectively stabilize each of the dimer structures. Additionally, we intend to form the dityrosine covalent linkages to see if we indeed stabilize a dimer with the structure covered in C1. Then, we will test the functional routes of these αSyn variants and by that gain better understanding as to their potential function.

## Acknowledgements

We would like to thank: (i) Drs. Asaf Grupi, Dan Amir, and Elisha Haas from the Mina & Everard Goodman Faculty of Life Sciences in Bar Ilan University for sharing the plasmids of αSyn bearing single cysteine mutations, (ii) Dr. Yuval Garini for inviting us to perform the bulk inter-molecular time-resolved- and FCS-FRET measurements of the mixtures of donor-and acceptor-labeled residues 39 αSyn on his experimental setup, (iii) Dr. William Breuer from the Proteomics and Mass Spectrometry Unit in the Alexander Silberman Institute of Life Sciences, the Hebrew University of Jerusalem, for assisting us with intact protein mass spectrometry, (iv) Drs. David Eliezer, Philipp Selenko, Hagen Hofmann, Yifat Miller, Deborah Toiber and Rina Rosenzweig for fruitful and insightful discussions. We acknowledge support from the National Institutes of Health (NIH) 1R35 GM134864 and the Passan Foundation (to N.V.D.). The project described was supported by the National Center for Advancing Translational Sciences, NIH, through grant UL1 TR002014. The content is solely the responsibility of the authors and does not necessarily represent the official views of the NIH (to N.V.D.). In addition, this project was supported by the Israel Science Foundation (grant 556/22 to E.L., grant 1768/15 to N.K.).

## Author contributions

J.Z. performed hetero-isotopic CL-MS experiments and analyzed their results, consulted by N.K., J.Z. performed western blot of crosslinked αSyn, J.Z., S.Z. and E.L. performed inter-molecular FRET and S.Z. performed time-resolved smPIFE experiments, with the assistance of P.D.H. in data analysis, S.Z., J.Z., P.D., M.L. and E.L. performed chromatography coupled to MALS experiments, J.C. and N.V.D. performed CL-MS-restrained DMD simulations. J.Z., S.Z., J.C., N.V.D. and E.L. wrote the paper and all co-authors assisted in refining it.

## Methods

### Expression and purification of recombinant αSyn variants

The expression and production of recombinant αSyn variants (*wt* and Cys mutants given to us as a gift; see Acknowledgements) is performed following previous existing protocols (Grupi & Haas, 2011; Zaer & Lerner, 2021). Briefly, αSyn is expressed from a pT7-7 plasmid encoding either the WT or the single Cys mutant proteins. Plasmids are transformed into BL21(DE3) component cells. The purification of the *wt* and mutant αSyn proteins are done by cells lysis, followed by osmotic shock and DNA precipitation, ending with proteins precipitation by ammonium sulfate. To further separate and purify the αSyn, the protein samples are loaded into 1 mL MonoQ column using FPLC system. The eluted fractions are loaded into 12% SDS-PAGE gel, relevant fractions, which exhibit a single band with molecular mass of αSyn (Fig. S19) are unified and dialyzed by 8 KDa MWCO dialysis bags at 4 °C against 30mM Tris, 2 mM EDTA buffer. The concentration was determined by spectrophotometer at 280 nm. Protein samples are aliquoted and stored immediately at -20 °C and aliquots were thawed shortly before the measurements.

### Chromatography coupled to MALS

AIEX- and SEC-MALS experiments are performed using the same setup as was previously reported (Amartely *et al*., 2018; Some *et al*., 2019). Briefly, we use a miniDAWN TREOS multi-angle light scattering detector, with three angle detectors (43.6°, 90°, and 136.4°) and a λ=658.9 nm laser beam (Wyatt Technology, Santa Barbara, CA) with a Wyatt QELS dynamic light scattering module for determination of hydrodynamic radius and an Optilab T-rEX refractometer (Wyatt Technology) set in-line with either a 1 mL Mono-Q analytical column (GE, Life Science, Marlborough, MA) for AIEX-MALS or a Superdex 75 or 200 column (GE, Life Science, Marlborough, MA) for SEC-MALS. Experiments are performed using an Äkta Pure M25 system with a UV-900 detector (GE) adapted for analytical runs. All AIEX-MALS experiments are performed at room temperature (25 °C) at 1.5 mL/min, using 30 mM Tris-HCl buffer pH = 8, as equilibration buffer, and 30 mM Tris-HCl buffer pH = 8 with 500 mM NaCl as elution buffer. 1 mg of αSyn at 0.2 mg/mL are injected to the column. Wash with 20 column volumes (CVs) of 0% elution buffer and 7 CVs of 30% elution buffer and eluted by 30 CVs gradient of 30-100% elution buffer collecting 1 mL fractions. Data collection and AIEX-MALS analysis are performed using the ASTRA 6.1 software (Wyatt Technology). The refractive index (RI) of the solvent is defined as 1.331, and the viscosity is defined as 0.8945 cP (common parameters for PBS buffer at λ=658.9 nm). dn/dc (the RI increment with protein concentration) value for all samples is defined as 0.185 mL/g (a standard average value for proteins). The molecular mass is calculated by both the absorption at λ=280 nm and RI. We use blank baseline reduction for calculations according to RI. We identify similar molecular mass values using the two types of calculations and present the molecular mass values calculated using the absorption at λ=280 nm.

### Cross-linking followed by western blot

*Wt*-αSyn was dialyzed by 8 KDa MWCO dialysis bags at 4 °C against HEPES buffer pH.7.5 and diluted to concentrations of 5 or 10 μM. We dissolve bis- sulfosuccinimidyl suberate (BS^3^) powder (Sigma Aldrich) in HEPES buffer (pH 7.18) to a concentration of 10 mM. We add the prepared BS^3^ to each of the αSyn mixtures to a final concentration of 1 mM and incubated at 4 °C for 1.5 h with shaking at 600 rpm. We quenched the cross-linking reaction by the addition of 20 mM ammonium bicarbonate for 20 min.

Following the cross-linking samples were resuspended in sample buffer, boiled at 96°C for 5 minutes, and centrifuged at 3,000 g for 30 seconds. Samples were then loaded onto a 12% acrylamide SDS-PAGE gel and transferred to 0.45 μM PVDF blotting membrane -Amersham HyBond (GE health care) using a Trans-blot Turbo System (BioRad, Hercules, CA), using a 1 hour program of 25 V, 2.5 A. The membrane was then treated with a blocking solution for half an hour (5% skim milk in 1% TBST) and incubated with the primary antibody (ab138501, Abcam, Cambridge, UK) at a ratio of 1:5,000 overnight at 4°C under constant motion. The membrane was then washed three times with TBST and incubated with the secondary antibody (111035144, Jackson ImmunoResearch, West Grove, Pennsylvania) at a ratio of 1:5,000 for 60 minutes at room temperature under constant motion. The membrane was then briefly washed again three times, incubated for ∼1 minute with ECL (ThermoScientific) under constant motion at room temperature, and photographed using a Lumitron photography system.

### αSyn dye labeling

αSyn dye labeling with either ATTO 488, ATTO 643, ATTO 647N (ATTO-TEC, GmbH) or sulfo-Cy3, linked to either iodoacetamide or maleimide thiol-reactive groups, are coupled specifically to single Cys residues in αSyn mutants via a thiol coupling reaction, as previously described (Zaer & Lerner, 2021).

### Steady-state FRET measurements

Acceptor excitation spectra are recorded using a spectrofluorometer (Jasco FP8200ST, Japan), scanning excitation wavelength in the range 400-650 nm and focusing on acceptor emission at λ=665 nm. Acceptor excitation spectra are normalized to the excitation spectrum part, in the wavelength range for direct acceptor excitation (550-650 nm; see Fig. S10). Then, the excitation spectra part in the wavelength range that covers donor excitation (440-550 nm; see Fig. S10) serves as a reporter for FRET.

### Time-resolved FRET and FCS-FRET measurements

FRET measurements of donor- and acceptor-labeling at residue 39 of αSyn are performed on a confocal fluorescence microscope (MicroTime200, PicoQuant GmbH Berlin, Germany) assembled on top of a modified Olympus IX71 inverted microscope stand, the same as previously reported (Lerner *et al*., 2013). Briefly, pulsed picosecond diode lasers are used for the excitation of the donor and acceptor (LDH-P-C-470B and LDH-P-C-640B, respectively, PicoQuant GmbH Berlin Germany) with output wavelengths of 470 and 640 nm, respectively (60 and 40 μW average-power at the sample, respectively). The conservation of polarization modes before reaching the sample is verified for both laser sources. Bandpass filters ensured that only light within the desired excitation band reached the sample. The repetition rate is set to 20 MHz. A dichroic beam splitter with high reflectivity at 470 and 635 nm reflects the light through the optical path to a high numerical aperture (NA) super Apo-chromatic objective (60X, NA=1.2, water immersion, Olympus, Japan), which focuses the light onto a small confocal volume. Fluorescence from the excited molecules is collected through the same objective and focused with an achromatic lens (f = 175 mm) onto a 50 μm diameter pinhole. The fluorescence emissions of the donor and the acceptor are separated using a dichroic long pass filter with the dividing edge at 605 nm (FF605-Di01, Semrock Rochester NY, USA). Emission that passes through the donor channel is selected with a 520/35 nm band-pass filter (FF01-520/35-25, Semrock Rochester NY, USA); the emission passed to the acceptor channel is selected by a 690/70 nm band-pass filter (HQ690/70, Chroma Technology Corp. Rockingham Vermont, USA). The detectors used are single-photon avalanche photodiodes (SPAD; Donor Channel: 100 μm, MPD PD1CTC, Acceptor Channel: 170 μm, Perkin Elmer SPCM-AQRH 13). The data is correlated by the HydraHarp400 time correlated single photon counting (TCSPC) system and collected with SymphoTime version 5 (both by PicoQuant GmbH Berlin, Germany). The fluorescence decay curves are recorded as TCSPC histograms with bin widths of 16 ps.

Similarly, time-resolved FRET measurements of a mixture of αSyn labeled at residue 39 and αSyn labeled at residue 140 were performed on a system that was reported recently (see (Harris *et al*., 2021)), only with different donor conditions: pulsed laser at 488 nm for donor excitation (instead of 532 nm), with dichroic mirror splitting the fluorescent signals at 605 nm and with a 510/20 nm bandpass filter.

Recorded data is stored in the time-tagged time-resolved (TTTR) format, allowing each detected photon to be recorded together with its individual timing relative to the beginning of the measurement, and relative to the time of the excitation pulse, as well as the detection channel. The presented measurements are performed at an approximate depth of 70 μm inside a drop of solution, enclosed between two glass coverslips, sealed by a silicon double sticky sheet (Sigma). The total acquisition times are several minutes.

### smPIFE measurements

All measurements are performed on the same setup and are analyzed exactly as recently explained (Zaer & Lerner, 2021), just in the presence of varying *wt*-αSyn concentrations.

### Data analysis for PIE-smFRET

Resulting acquisition files were converted into the universal photon HDF5 file format using the Python code phconvert (Ingargiola, Laurence, *et al*., 2016a). Then, single molecule burst analysis was carried out as described previously (Ingargiola *et al*., 2018; Lerner *et al*., 2018) using the FRETbursts Python-based software g(Ingargiola, Lerner, *et al*., 2016). For selecting all bursts (Fig. S18, A), we used a burst selection threshold of 50 photons in a burst. For selecting only bursts exhibiting fluorescence arising from donor excitation and from acceptor excitation (Fig. S18, B), we used a burst threshold of 25 photons from the donor excitation period and 25 photons from the acceptor excitation period.

### Intact protein mass determination

We characterize the molar mass of the purified αSyn by intact protein mass spectrometry and the results report on the exact molecular mass of the αSyn (Fig. S20). The procedures is hereby described: Proteins are dissolved in 40% acetonitrile, 0.3% formic acid (all solvents were MS-grade) at a concentration of 2- 5 mg/mL. Then, the dissolved proteins are injected directly via a HESI-II ion source into a Q Exactive Plus (Thermo Fisher Scientific) mass spectrometer and a minimum of three scans lasting 30 seconds are obtained. The scan parameters are: i) scan range 1,800 to 3,000 m/z without fragmentation; ii) resolution 140,000; iii) positive polarity; iv) AGC target 3×10^6^; v) maximum inject time 50 ms; vi) spray voltage 4.0 kV; vii) capillary temperature 275 °C; and viii) S-lens RF level 55. Scan deconvolution is done using Mag Tran version 1.0.3.0 (Amgen).

### Hetero-isotopic CL-MS

#### Testing the ^15^N coverage in ^15^N-αSyn

We digest ^14^N- and ^15^N-αSyn (Alexo Tech, Umea, Sweden) separately and observe the peptides. We examine the spectrum of a specific peptide’s mass in the ^15^N-αSyn and find that the mass precisely equals the mass of ^14^N-αSyn of the exact same peptide plus ∼1 Da multiplied the number Nitrogen atoms present in the peptide.

If the labeling is not full, we will obtain overlapping between the series of the peaks with a difference in the mass that is corresponding to the number of nitrogen atoms that are not labeled in the peptide. The analysis takes into consideration the ^13^C isotope series and in each peptide, the expected mono-isotopic mass of the examined peptides is the first peak in the isotopic series without preceding peaks that could be unlabeled nitrogen atoms.

#### Cross-linking of α-synuclein with BS3

Equimolar ratio mixtures of ^14^N, ^15^N (AlexoTech) or ^14^N-^15^N αSyn are prepared to a final concentration of 2.5 μM. We dissolve bis-sulfosuccinimidyl suberate (BS^3^) powder (Sigma Aldrich) in HEPES buffer (pH 7.18) to a concentration of 10 mM. We add the prepared BS^3^ to each of the αSyn mixtures to a final concentration of 1 mM and incubated at 4 °C for 1.5 h with shaking at 600 rpm. We quenched the cross-linking reaction by the addition of 20 mM ammonium bicarbonate for 20 min (Kalisman *et al*., 2012; Tayri-Wilk *et al*., 2020).

#### Cross-linking of α-synuclein with DMTMM

Equimolar ratios mixtures of ^14^N, ^15^N (AlexoTech) or ^14^N-^15^N αSyn are prepared to a final concentration of 2.5 μM. We dissolve 4-(4,6-dimethoxy-1,3,5-triazin-2-yl)-4-methyl-morpholinium chloride (DMTMM) powder (Sigma Aldrich) in HEPES buffer (pH 7.18) to a concentration of 70 mM. We add the prepared DMTMM to each of the αSyn mixtures to a final concentration of 7 mM and incubated at 4 °C for 1.5 h with shaking at 600 rpm. We quenched the cross-linking reaction by the addition of 20 mM ammonium bicarbonate for 20 min (Kalisman *et al*., 2012; Tayri-Wilk *et al*., 2020).

#### Preparation of samples for mass spectrometry

The proteins are precipitated in 1 mL of acetone (-80 °C) for one hour, followed by centrifugation at 14,000 g. The pellet is resuspended in 20 μL of 8 M urea. The urea is diluted by adding 200 μL of digestion buffer (25 mM TRIS, pH=8.0; 10% Acetonitrile). We add 0.5 μg of trypsin (Promega, Madison, Wisconsin) or 0.5 μg GluC (Promega) for the DMTMM to the diluted urea and digest the protein overnight at 37°C under agitation. Following digestion, the peptides are desalted on C18 stage-tips and eluted by 55% acetonitrile. The eluted peptides are dried in a SpeedVac, reconstituted in 0.1% formic acid, and measured in the mass spectrometer (Kaledhonkar *et al*., 2018).

#### Mass spectrometry

The samples are analyzed by a 120 minute 0-40% acetonitrile gradient on a liquid chromatography system coupled to a Q-Exactive HF mass spectrometer. The analytical column is an EasySpray 25 cm heated to 40°C. The method parameters of the run are: i) Data-Dependent Acquisition; ii) Full MS resolution 70,000 ; iii) MS1 AGC target 1e6 ; iv) MS1 Maximum IT 200 ms ; v) Scan range 450 to 1800 ; vi) dd-MS/MS resolution 35,000 ; vii) MS/MS AGC target 2e5 ; viii) MS2 Maximum IT 600 ms ; ix) Loop count Top 12; x) Isolation window 1.1 ; xi) Fixed first mass 130 ; xii) MS2 Minimum AGC target 800 ; xiii) Peptide match - off ; xiv) Exclude isotope - on ; and xv) Dynamic exclusion 45 seconds. Each cross-linked sample is measured twice in two different HCD energies (NCE): 26, and stepped 25, 30, and 35. All cross-linked samples are measured with the following charge exclusion: unassigned,1,2,3,8,>8. Proteomics samples are measured with the following charge exclusion: unassigned,1,8,>8.

#### Identification of cross-links

The RAW data files from the mass spectrometer are converted to MGF format by Proteome Discoverer (Thermo). The identification of BS^3^ or DMTMM cross-links uses a search application (FindXL: http://biolchem.huji.ac.il/nirka/software.html) that exhaustively enumerates all the possible peptide pairs. The search parameters are as follows: i) Sequence database- ^14^N α-synuclein and ^15^N α-synuclein that differs in the molecular weight of the amino acids by 1 Da multiplied by the number of Nitrogen in the amino acid; ii) Protease – trypsin or GluC, allowing up to three mis-cleavage sites; iii) Variable modifications: methionine oxidation, lysine with hydrolyzed mono-link; iv) Cross-linking must occur between two lysine residues; v) Cross-linker is not cleaved; vi) MS/MS fragments to consider: b-ions, y-ions; vii) MS1 tolerance – 6 ppm; viii) MS2 tolerance – 8 ppm; and ix) Cross- linker mass – one of three possible masses: 138.0681, 138.0681 + 1.00335, and 138.0681 + 2.0067 for the BS^3^ and -18 Da for the DMTMM that represent the release of water molecule. The three masses address the occasional incorrect assignment of the mono-isotopic mass by the mass spectrometer.

### Restrained DMD simulations

#### Preparation of restraints for DMD simulations from CL-MS results

BS^3^ and DMTMM lead to the cross-linking of specific pairs of amino acids and cover these pairs if they were in a structure with Cα-Cα distances as long as 30 and 16 Å, respectively. To maximize the efficiency of using these results in structural modeling, we used the Cα-Cα distance of DMTMM, to design DMTMM-related discrete potential functions as restraints, however for BS^3^, we chose instead the inter-residue end-to-end distance to be within the distances covered by the length of the cross-linker (max. 11.1 Å). Unlike in rigid protein complexes, where the cross-linking data can be used as if only these distances occur within the complex, in a protein as flexible and dynamic as αSyn, inter-residue distances longer than the range covered by the cross-link are possible. We took this into account when designing discrete restraining potentials for DMD simulations, by allowing all inter-residue distances to occur, but with an energetic preference towards the range covered by the cross-linker (Fig. S11).

#### CL-MS-restrained DMD simulations and validations

Cross-linking-restrained discrete molecular dynamics (DMD) simulations were developed to solve protein structures successfully (Brodie *et al*., 2017, 2019). Based on the same strategy, we apply 121 intra-molecular and 36 inter-molecular cross-links to restrain DMD simulations of the αSyn dimer. Intra-molecular cross- links restrain the distance between residues within a single subunit, while inter- molecular cross-links restrain the distance between residues from two different subunits. We convert the cross-link distance ranges to square-well potential functions (Fig.S11), which are incorporated with the Medusa force field during simulations (Ding & Dokholyan, 2006; Dokholyan, 2006; Proctor *et al*., 2011; Shirvanyants *et al*., 2012; J. Wang & Dokholyan, 2019; Yin *et al*., 2008). We start simulations by using two fully extended conformations of αSyn that are separate from each other. We conduct an all-atom replica exchange DMD simulation where 26 replicas with temperatures ranging from 0.400 to 0.800 kcal/(mol•kB) are run for 4 x 10^6^ time steps (50 ps per time step) (Ding *et al*., 2008).

After simulations, we validate the resulting dimer structure models by comparing them with CL-MS and smPIFE experimental data. We calculate the distances between cross-linked residues within all the simulated dimer structures using GROMACS and plot the distance distribution histogram for each cross-link. If the maximum length of the cross-linker falls within or above the distance distribution that indicates this cross-link is satisfied by our simulation. On the other hand, if the maximum length is smaller than the overall distance distribution that means this cross-link is not captured by our simulation results. Moreover, we calculate the percentage of structures within each cluster that are validated by cross-linking constraints. To compare our dimer structure models with smPIFE data, we calculate the accessible volumes of residues 26, 39, 56, and 140 within each monomer for each one of the eight clusters using Python libraries MDTraj (https://www.mdtraj.org/1.9.5/index.html) and AvTraj (https://github.com/Fluorescence-Tools/avtraj). Then, we measure the solvent accessible surface area (SASA) of each volume using PyMOL (https://pymol.org/2/#download). The sub-populations indicated by the surface area of all the structures within each cluster are compared with the number of sub- populations identified by smPIFE experiments.

#### Analysis of αSyn dimer structure models

We extract all the dimer structure models from the 26 trajectories. We calculate the radius of gyration (Rg) of all the dimers and plot Rg versus energy. We perform agglomerative hierarchical clustering of the most populated structures using the ward linkage algorithm from TTClust (Tubiana *et al*., 2018). We optimize the clustering cutoff until the average RMSD values between clusters are much larger than the average RMSD values within each cluster (Tables S2 and S3). We calculate the average SASA of each residue for all the clusters using the GROMACS SASA algorithm (Eisenhaber *et al*., 1995). The SASA values are further divided by the maximum surface area of the corresponding residue to generate normalized SASA. A threshold of 40% is applied to determine the buried and exposed states (Stultz *et al*., 1993; Ullman *et al*., 2011).

## Data Availability

The raw photon-HDF5 (Ingargiola, Laurence, *et al*., 2016b) files of all smPIFE measurements, and the data analysis pipeline using the FRETbursts (Ingargiola, Lerner, *et al*., 2016) software, are available publicly as Jupyter Notebooks over Zenodo (Lerner *et al*., 2021b).

The mass spectrometry proteomics data have been deposited to the ProteomeXchange (Deutsch *et al*., 2020) Consortium via the PRIDE (Perez-Riverol *et al*., 2019) partner repository with the dataset identifier PRIDE: PXD030299.

The models of the aSyn ensemble structure have been deposited in the PDB-Dev database (Vallat *et al*., 2018) with the accession number PDB-Dev: PDBDEV_00000098.

