## Supplementary material for "Structural and Dynamic Insights Into α-Synuclein Dimer Conformations": SI

#### Supporting Information

Joanna Zamel[1,*], Jiaxing Chen[2,*], Sofia Zaer[1], Paul David Harris[1], Paz Drori[1], Mario Lebendiker[3], Nir Kalisman[1], Nikolay V. Dokholyan[2,4,5,#], Eitan Lerner[1,6,#]

[1] Department of Biological Chemistry, The Alexander Silberman Institute of Life Sciences, Faculty of Mathematics & Science, The Edmond J. Safra Campus, The Hebrew University of Jerusalem, Jerusalem 9190401, Israel

[2] Department of Pharmacology, Penn State College of Medicine, Hershey, PA 17033, USA

[3] Wolfson Centre for Applied Structural Biology, The Alexander Silberman Institute of Life Sciences, The Hebrew University of Jerusalem, The Edmond J. Safra Campus, Givat Ram, Jerusalem, 9190401, Israel

[4] Department of Biochemistry & Molecular Biology, Penn State College of Medicine, Hershey, PA 17033, USA

[5] Departments of Chemistry and Biomedical Engineering, Pennsylvania State University, University Park, PA 16802, USA

[6] The Center for Nanoscience and Nanotechnology, The Hebrew University of Jerusalem, Jerusalem 9190401, Israel

[*] Shared first authorship

[#] Corresponding authors

#### Details of additional fluorescence-based experiments

##### Steady-state inter-molecular FRET: small-scale self-association equilibrium

Freshly-thawed αSyn in solution (see *Methods*) should exhibit mainly the monomeric form and if any oligomeric form exists in the sample, they should yield a minimal inter-molecular FRET signature. We measure the acceptor (ATTO 647N) excitation spectrum of a mixture of 100 nM donor (ATTO 488)-labeled αSyn molecules mixed with 100 nM acceptor-labeled αSyn molecules focusing on the acceptor fluorescence intensity (see *Methods*). Therefore, FRET reports uniquely on inter-molecular interactions. As a control, we measure the acceptor excitation spectrum of a sample containing 100 nM of only acceptor-labeled αSyn, to account for all the non-FRET spectral signatures, hence solely on direct acceptor excitation.

Our results exhibit a peak in the acceptor excitation spectrum at λ~500 nm (Fig. S5), which reports on the occurrence of inter-molecular FRET. The FRET signal confirms the presence of self-associated αSyn species, which combined with our MALS and WB-based evidence are primarily dimers.

##### Fluorescence correlation spectroscopy inter-molecular FRET: small-scale self-association equilibrium

To further test the abundance of the αSyn self-associated species at lower concentrations, we perform fluorescence correlation spectroscopy (Elson & Magde, 1974) (FCS)-FRET (FCS-FRET) measurements. These experiments measure the donor (ATTO 488) and acceptor (ATTO 647N) fluorescence fluctuations of freely-diffusing donor-labeled αSyn and acceptor-labeled αSyn in decreasing concentrations. To reduce the possibility for multiple labeled molecules to enter the confocal volume, we work with ever-decreasing total concentrations (e.g., 50, 12, 3 nM), and then calculate the donor fluorescence autocorrelation and the donor-acceptor fluorescence cross-correlation curves. If two or more monomers are in a sufficient proximity with each other while diffusing through the confocal spot of our system, then exciting the donor will lead to the energy transfer to the nearby acceptor and photons will be detected on both the donor and acceptor fluorescence detection channels. The donor-acceptor fluorescence cross-correlation curve reports on such species that traverse the effective excitation volume. If αSyn is in its monomeric state, donor fluorescence will mostly be detected. The donor fluorescence autocorrelation curve reports on both donor-labeled species: monomers and self-associated species. The ratio of the amplitudes of the donor-acceptor fluorescence cross-correlation and the donor fluorescence autocorrelation curves is proportional to the fraction of self-associated αSyn species labeled with both donor and acceptor (Buschmann et al., 2014). The results reported in Fig. S8 exhibit a fraction of donor- and acceptor-labeled αSyn self-associated species, labeled at residue 39, exists at αSyn concentrations as low as 3 nM. Overall, bulk steady-state-FRET, time-resolved FRET and FCS-FRET results point to an equilibrium between monomers and self-associated species of αSyn. Taken together with the results from MALS, this is a result of a mixture between primarily monomers and dimers of αSyn.

#### smPIFE: αSyn dimer formation involves local structural changes

To investigate the structural changes associated with the formation of an αSyn dimer, we perform sulfo-Cy3 (sCy3)-based intra-molecular time-resolved smPIFE measurements (Chen, Zaer et al., 2021; Zaer & Lerner, 2021). The uniqueness of this dye is that it exhibits excited-state isomerization between a bright *trans* isomer and a dark *cis* isomer, leading to an overall low fluorescence quantum yield. However, if the excited-state isomerization of sCy3 is obstructed due to steric hindrance, for instance by a protein surface in the vicinity of the dye, then the isomerization rate will decrease leading to more fluorescence from the *trans* isomer, and hence to a higher fluorescence quantum yield.

We hypothesize that like in the case of the free-form monomer (Chen, Zaer et al., 2021; Zaer & Lerner, 2021) the αSyn dimer may exhibit distinct sub-populations with different characteristic fluorescence lifetimes. To gain insights on the dimer formation we performed smPIFE measurements. We measured 25 pM of freely-diffusing sCy3-labeled αSyn in the presence of increasing concentrations of *wt*-αSyn, up to few μM to induce the formation of dimers (Fig. S9). Additionally, fitting of the mean nanotime histograms to a sum of Gaussian model may assist in the quantification of the mean fluorescence lifetimes and fractions of each sub-population (Table S1). From observing how the burst histogram of mean nanotimes change with increasing the *wt*-αSyn concentration, it is clear that: (i) sCy3 senses the local structural reorganization in the vicinity of residues 56 and 140, upon dimer formation, with an enhanced low mean nanotime sub-population, hence with minimal steric obstruction pointing towards more solvent exposure of the residues, (ii) sCy3 senses the local structural reorganization in the vicinity of residues 39, upon dimer formation, with both a low and intermediate mean nanotime sub-population, hence with steric obstruction levels pointing towards high and intermediate levels of solvent exposure, (iii) sCy3 fails to sense the local structural reorganization in the vicinity of residue 26, upon dimer formation, and (iv) some local structural changes when shifting from the monomer to the dimer include gradual changes at multiple *wt*-αSyn concentration concentrations and not necessarily a single transition. Importantly, in all sCy3-labeled residues, the results with the highest dimer fraction, at the highest *wt*-αSyn concentration exhibit dynamic heterogeneity between multiple mean nanotime sub-populations, with transitions between them slower than diffusion times through the confocal spot, hence slower than a few ms.

#### Supporting Figures

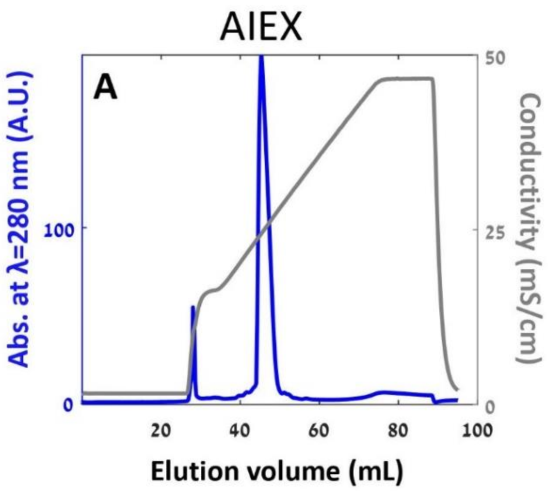


**Fig. S1: AIEX of wt-αSyn** using 1 mL mono Q column with the gradient shown in grey, has been performed on a sample of the WT αSyn. The sample has eluted as a single elution peak (absorption at a wavelength of 280 nm, blue).

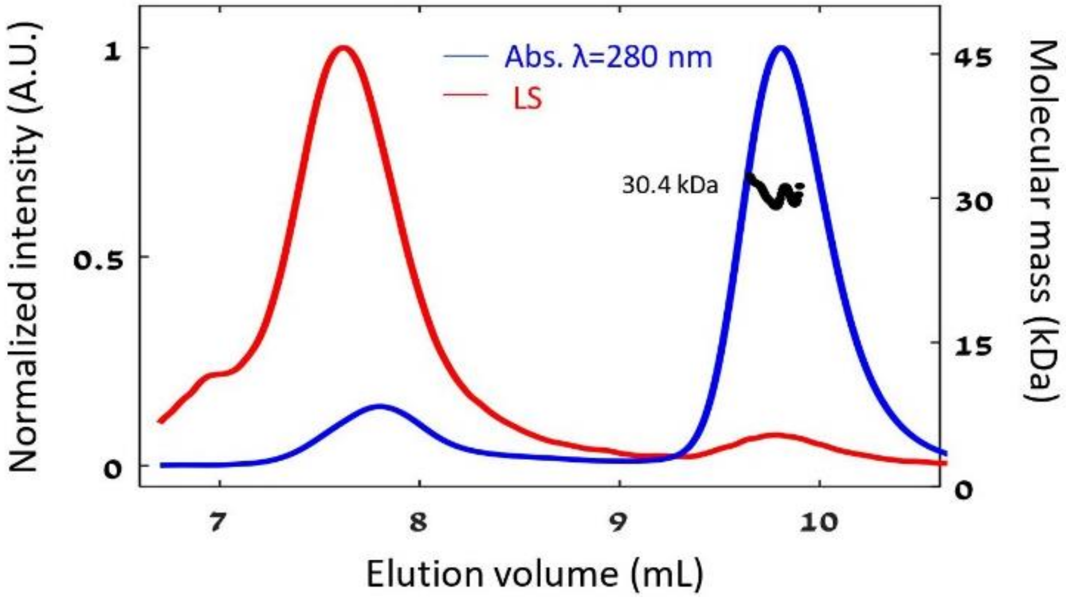


**Fig. S2: SEC-MALS of WT αSyn using Superdex75 column of *wt*-αSyn does not fully separate effect of aggregates, as does Superdex200 column.**

The late elution peak was characteristic of a species larger than a monomer, due to the MALS-derived molecular mass of 30.4±0.9 kDa that is larger than the monomer molecular mass of 14.4 kDa. The separation between the aggregates' elution peak and the monomer-dimer mixture elution peak was not good enough, which led to an increase in the average molecular mass of the main monomer-dimer mixture elution peak. Therefore, we moved to SEC-MALS with a Superdex200 column, which did not yield this overlap (see Fig. 1B).

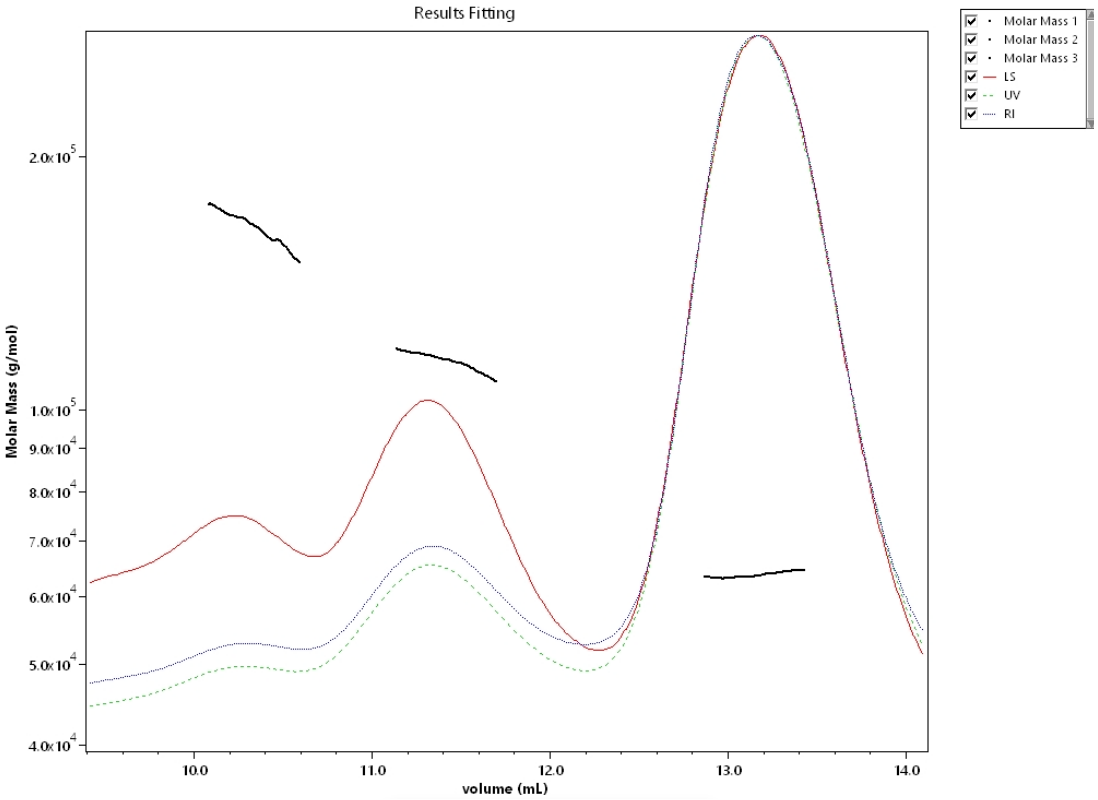


**Fig.S3: Calibration of Superdex200 column MALS using BSA**.
The light scattering intensity (red), protein absorption at 280 nm (green dashes) and refractive index, RI (blue) were monitored. The eluting peaks resulted in molecular masses of 164.20±1.43, 114.30±0.09 and 63.69±0.46 KDa, respectively, indicating trimer, dimer and monomer of BSA.

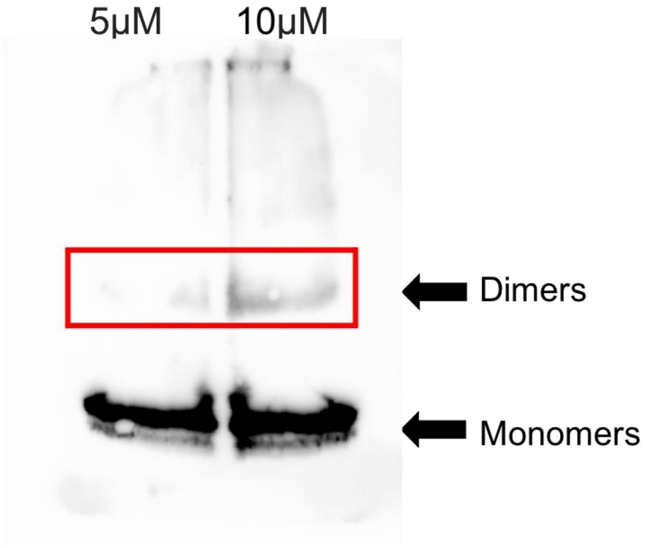


**Fig. S4: WB of BS**$^{3}$ **cross-linked *wt*-αSyn** in two different concentrations after 40 seconds exposure.

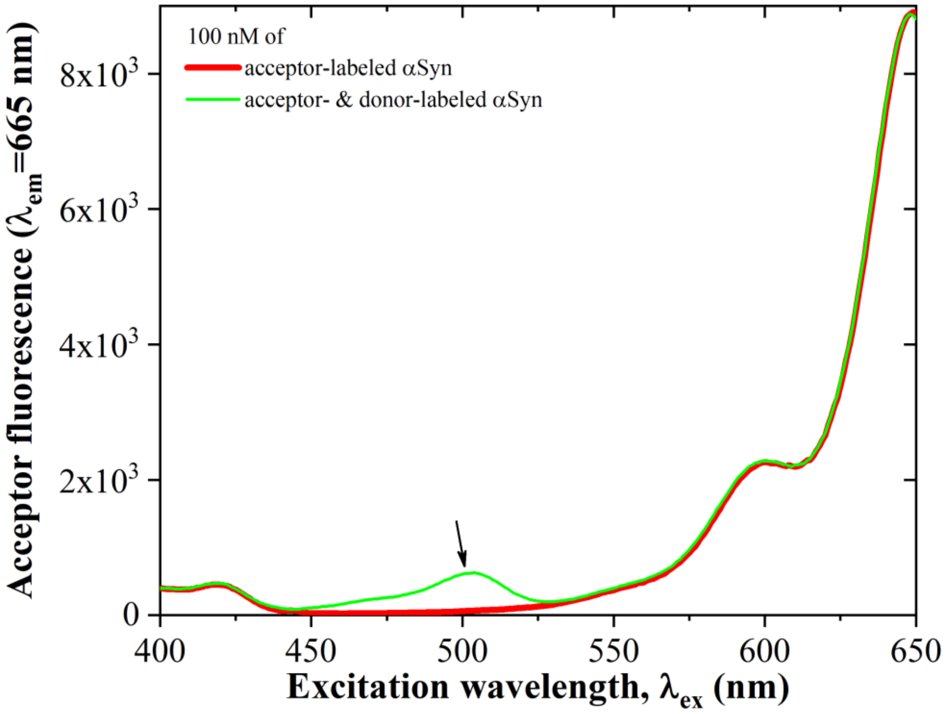


**Fig. S5 Acceptor excitation spectra show a FRET signature of dimers and short oligomers at 100 nM.** The acceptor excitation spectrum (focusing on emission at a wavelength of 665 nm) of a 100 nM mixture of donor- and acceptor-labeled at position αSyn was recorded on a freshly-thawed sample (green). The acceptor excitation spectra of 100 nM acceptor-labeled αSyn was also recorded (red). The excitation spectra shown were normalized to the excitation peak at the wavelength range in which direct acceptor excitation (Direct acceptor excitation) occurs (λ={525-650} nm). Then the excitation peak at the wavelength range λ={440-525} nm is indicative of inter-molecular FRET, that is a consequence of self-associated αSyn species.

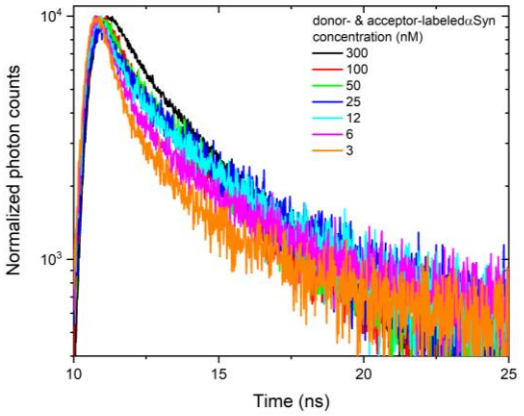


**Fig. S6: A signature of intermolecular FRET in acceptor fluorescence decay, decreases with αSyn concentration.**

The acceptor fluorescence decays, following donor excitation, were measured in position 39 donor- and acceptor-labeled αSyn mixtures at different nominal αSyn concentration (see inset). The fluorescence decay curves exhibit two components: a fast component, that is most probably due to leakage of the donor fluorescence red edge into the acceptor detection channel (see Fig. S7) and a slow component, indicative of acceptor excitations that were delayed in the donor excited state prior to the FRET event. The amplitude of the slow component is decreasing as the nominal αSyn concentration decreases.

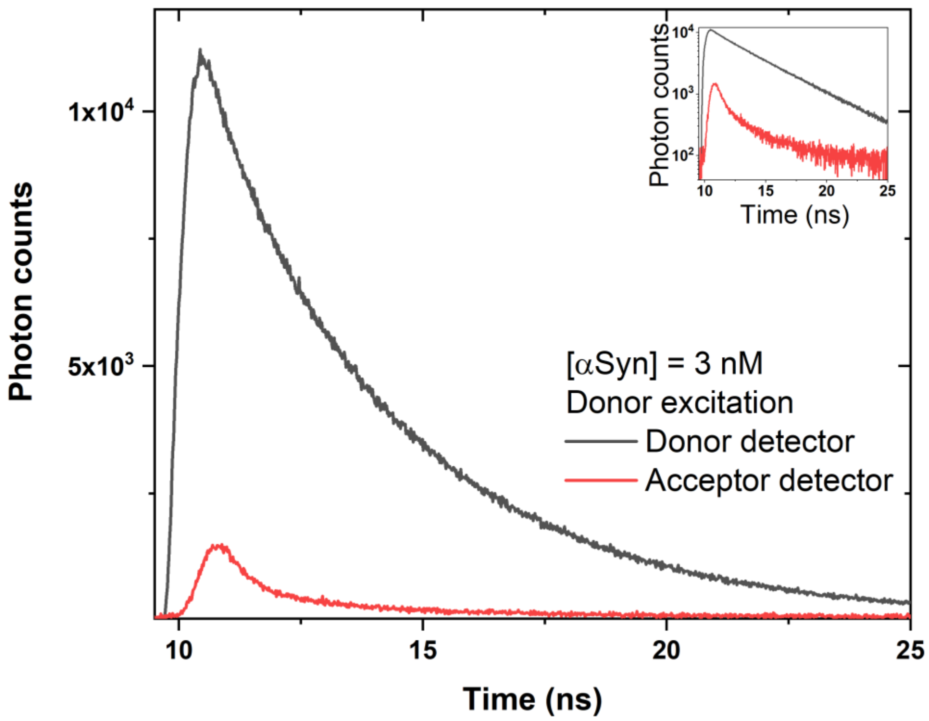


**Fig. S7: The signals recorded at the acceptor detection channel, when FRET does not occur, are due to direct acceptor excitation with at wavelength intended for donor excitation and donor fluorescence leakage into the acceptor detection channel.**
A fraction of the donor fluorescence decay (black) and other signal contaminations, such as direct acceptor excitation at the donor excitation wavelength, leak into the acceptor detection channel (red), yielding a minimal fluorescence lifetime in the acceptor detection channel. While the main panel reports the results in linear scale, the inset reports the results in log-scale.

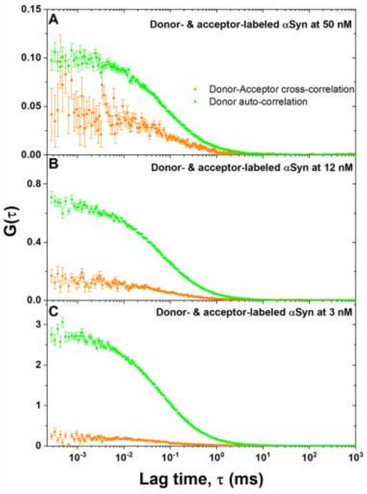


**Fig. S8**: **The abundance of a self-associated species of αSyn is decreasing with decreasing αSyn concentration.** Donor fluorescence auto-correlation curves (green) and donor-acceptor fluorescence cross-correlation curves (orange) of donor- and acceptor-labeled αSyn are shown at decreasing nominal concentrations (**A** - 50 nM; **b** – 12 nM; **c** – 3 nM). The amplitude of the donor-acceptor fluorescence cross-correlation relative to the amplitude of the donor fluorescence auto-correlation decreases with decreasing αSyn concentration, indicative of a decreasing concentration of species having coincidence of both donor and acceptor fluorescence (namely self-associated species).

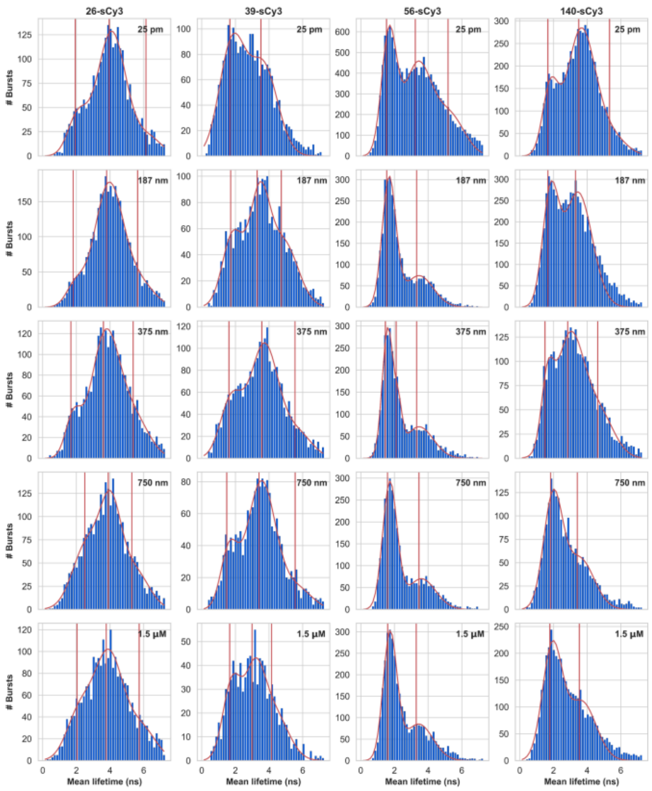


**Fig. S9**: **smPIFE data exhibiting how different sCy3-labeled residues sense a local structural change when changing from a monomer to a dimer.**

Mean fluorescence lifetime histograms of sCy3 labeling different residue positions (26, 39, 56, 140; columns) with 25 pM sCy3-labeled αSyn, in the presence of increasing concentrations (0, 187, 375, 750, 1,500 nM; rows) of unlabeled *wt*-αSyn. The concentrations shown at the top of panels signify the approximate overall αSyn concentration.

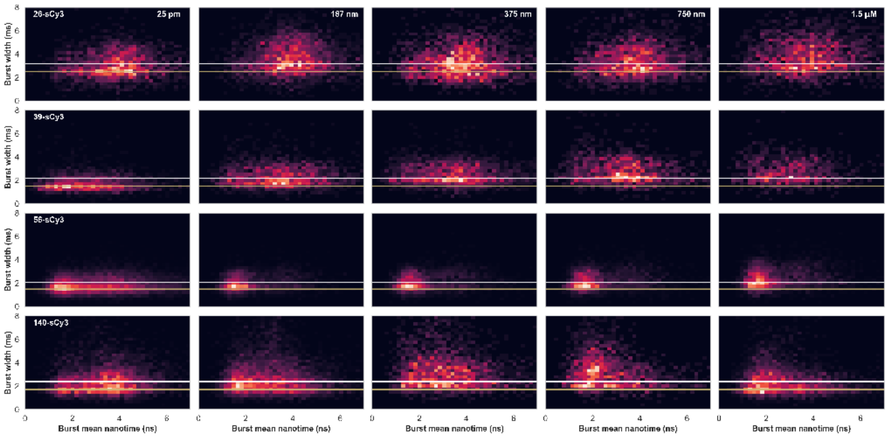


**Fig. S10: Increasing *wt*-αSyn concentration, in the presence of a constant 25 pM concentration of sCy3-labeled αSyn, leads to sCy3 fluorescence lifetime changes when labeling some residues and accompanied by a 30-50% increase in burst durations.**

Mean fluorescence lifetime (horizontal) and burst width (vertical) two-dimensional histograms of sCy3 labeling different residue positions (26, 39, 56, 140; rows) with 25 pM sCy3-labeled αSyn, in the presence of increasing concentrations (0, 187, 375, 750, 1,500 nM; columns) of unlabeled *wt*-αSyn. Yellow and White horizontal lines, running over all panel rows signify the peak values of two burst width subpopulations per each sCy3-labeled αSyn variant.

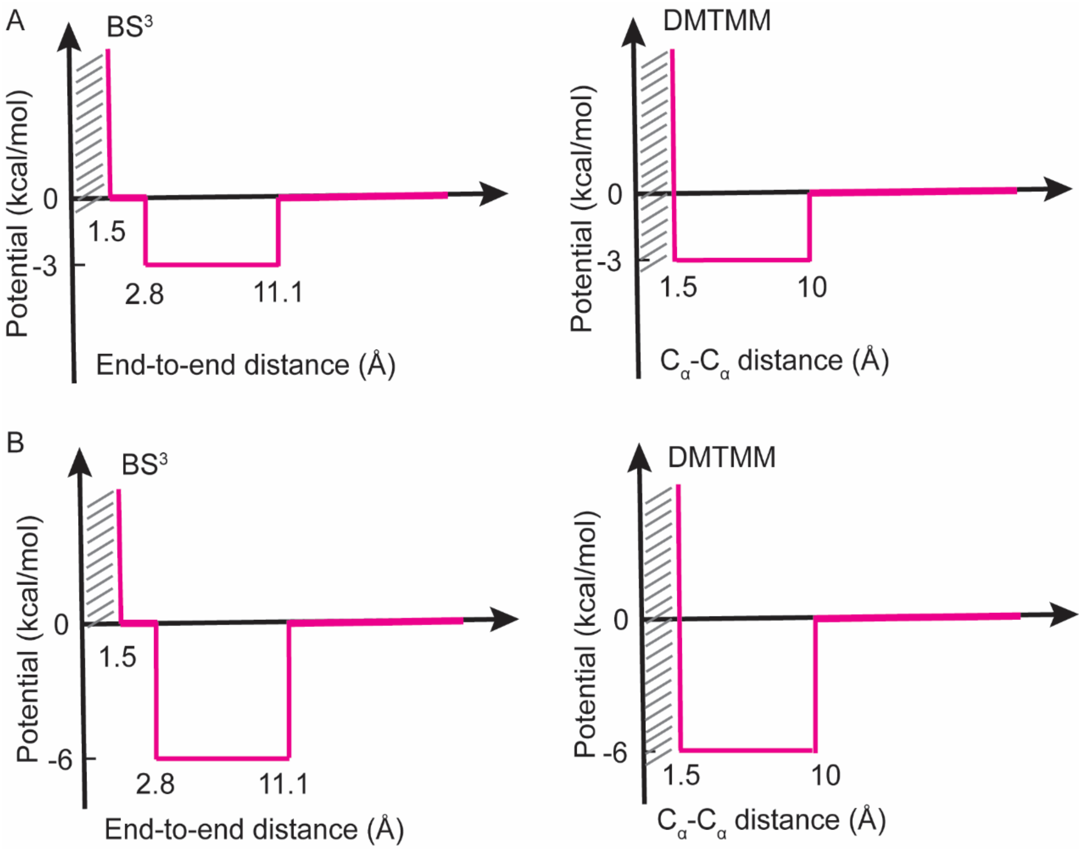


**Fig. S11: Design of potential functions acting as CL-MS-derived restraints for DMD simulations: intra-molecular (A) and inter-molecular (B) cross-links.**

While the $BS^3$ cross-links are used for restraining the end-to-end distances between the end atoms of the residue side chains, the DMTMM cross-links are used for restraining the $C_\alpha$-$C_\alpha$ distances. In these restraining potential functions, all distance values are possible, with an energetic preference towards the distance range covered by the cross-links. Note that inter-molecular cross-links have a higher energetic preference than do intra-molecular cross-links, to account for the fact that inter-molecular cross-links report on proximities between residues of different subunits in the dimer, and intra-molecular cross-links report intra-molecular proximities either in a dimer subunit or in a monomer that might exist in equilibrium.

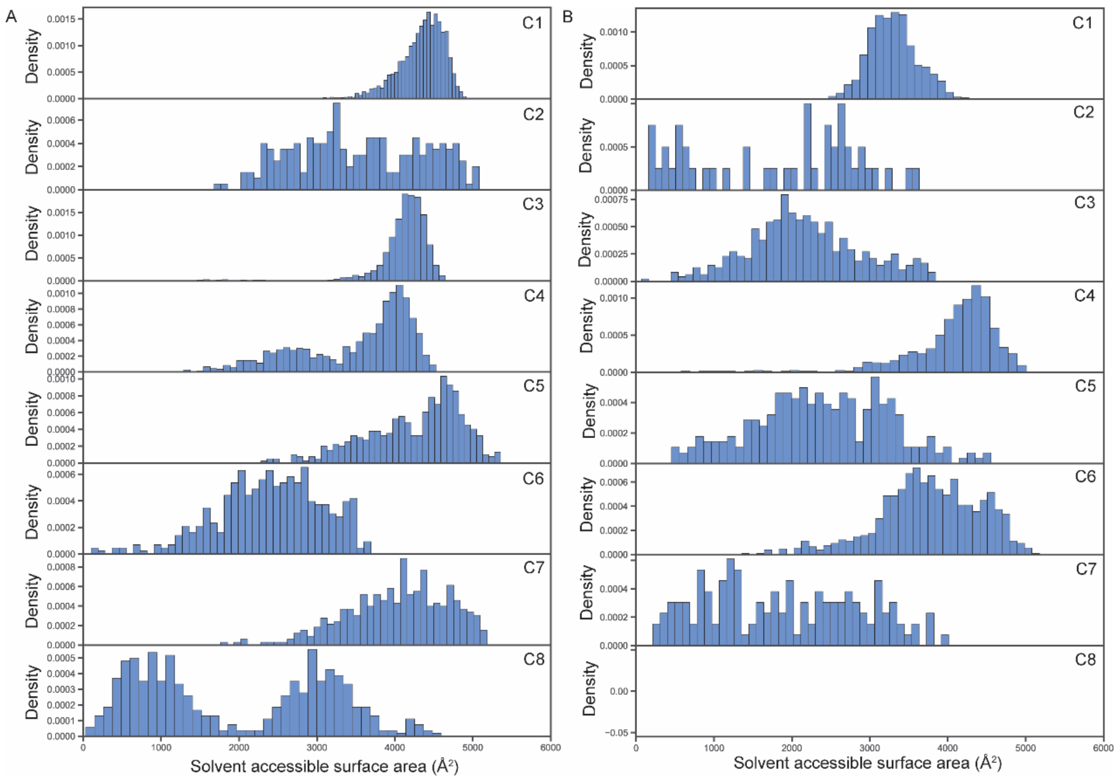


**Fig. S12: The distribution of solvent accessible surface area of the accessible volume of Cy3 dye conjugated with residue 26** in chain A (**A**) and chain B (**B**) of the dimer. C1-8 represent clusters 1-8.

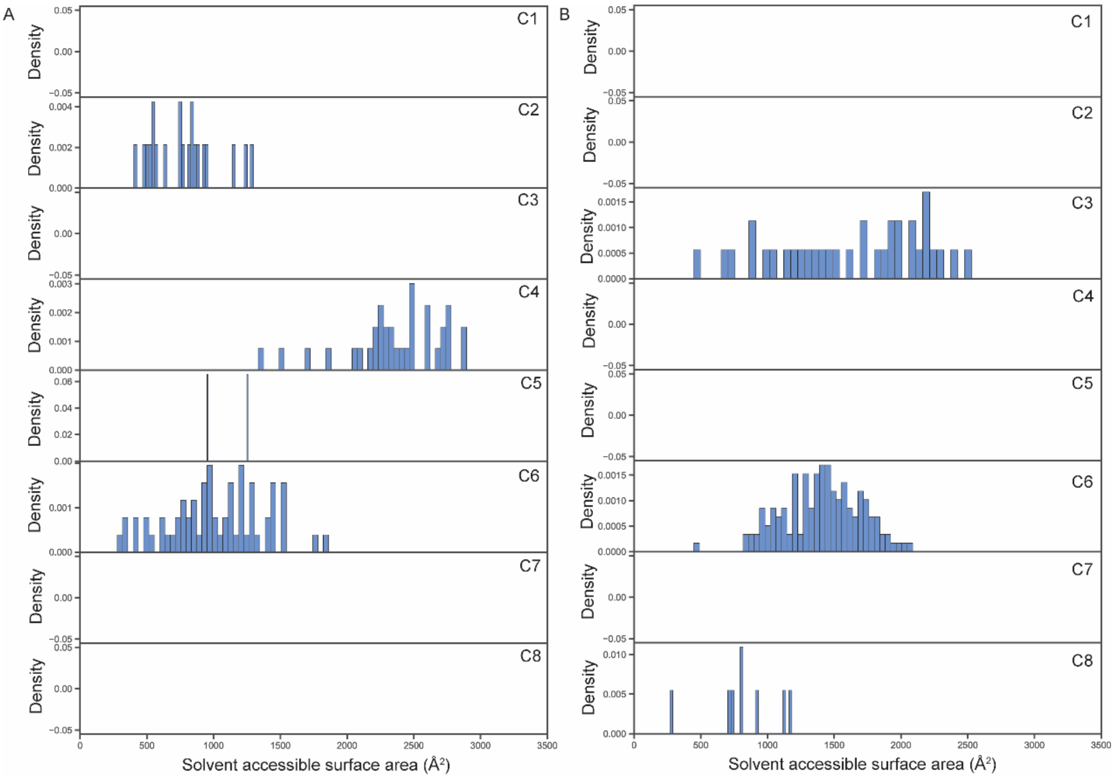


**Fig. S13: The distribution of solvent accessible surface area of the accessible volume of Cy3 dye conjugated with residue 39** in chain A (**A**) and chain B (**B**) of the dimer. C1-8 represent clusters 1-8.

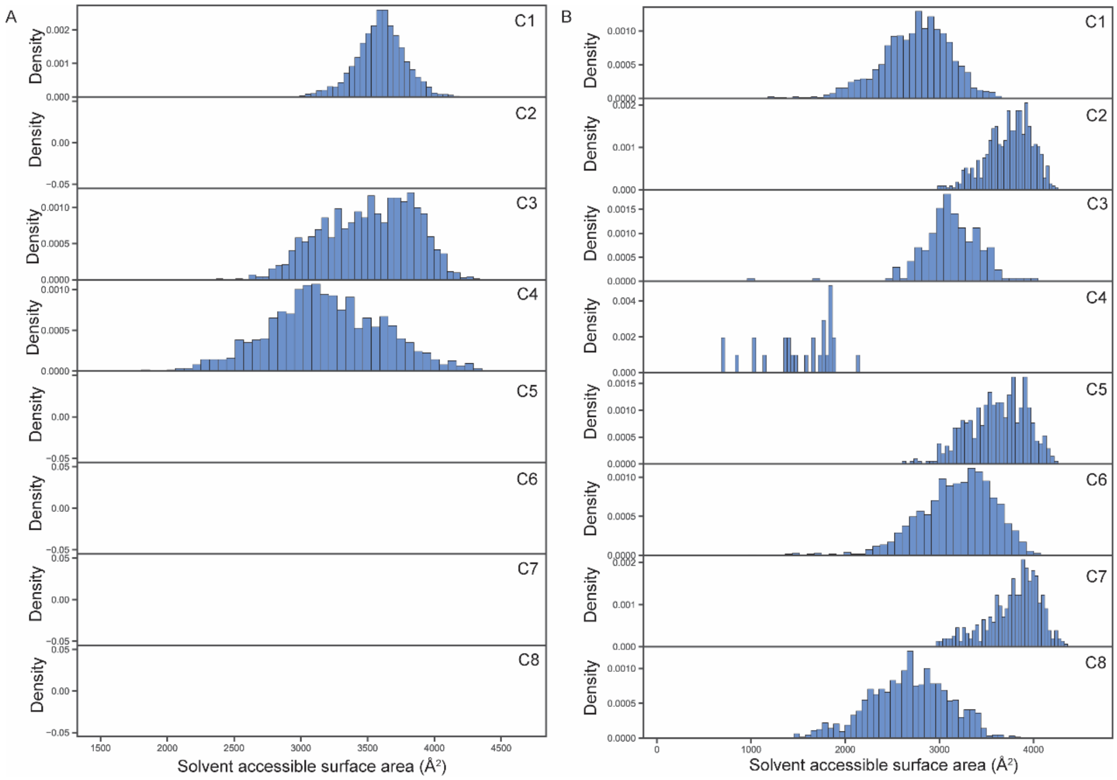


**Fig. S14: The distribution of solvent accessible surface area of the accessible volume of Cy3 dye conjugated with residue 56** in chain A (**A**) and chain B (**B**) of the dimer. C1-8 represent clusters 1-8.

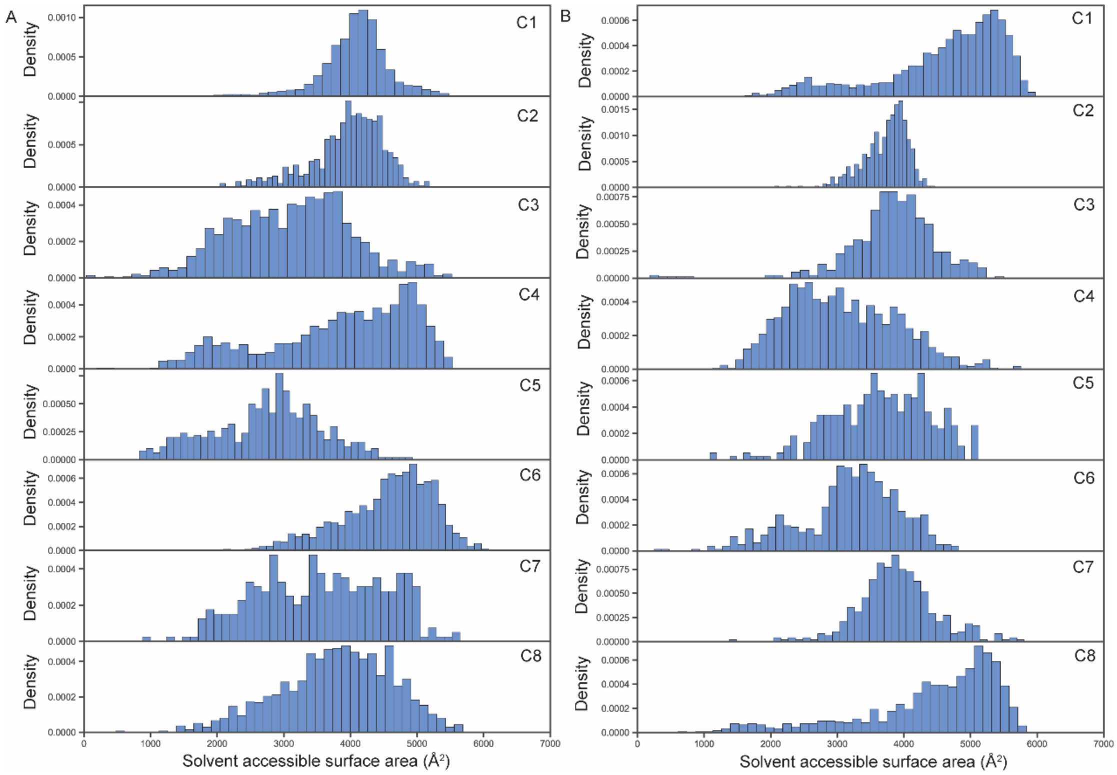


**Fig. S15: The distribution of solvent accessible surface area of the accessible volume of Cy3 dye conjugated with residue 140** in chain A (**A**) and chain B (**B**) of the dimer. C1-8 represent clusters 1-8.

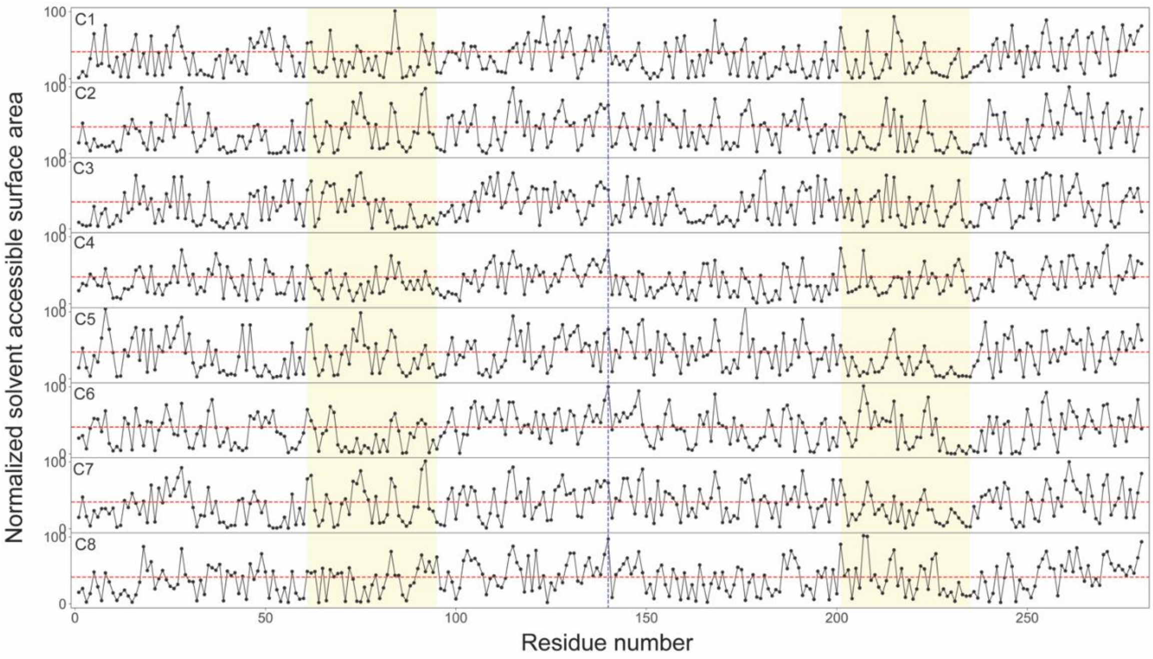


**Fig. S16: The normalized solvent accessible surface area of each residue for the 8 clusters.** Residue numbers 1-140 indicate chain A and residue numbers 141-280 represent chain B. C1-8 represent clusters 1-8.

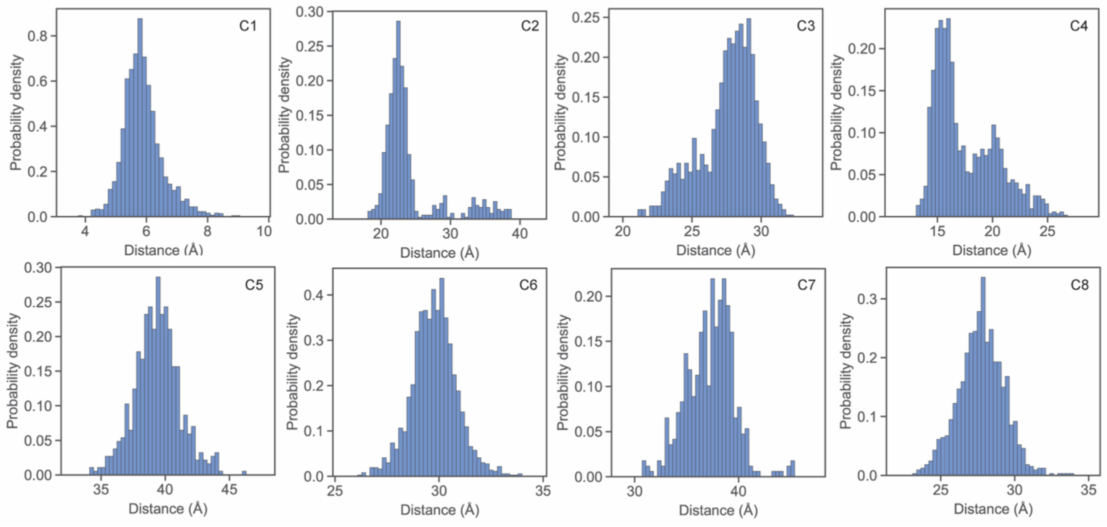


**Fig. S17: The distance distribution between residues 39 of each monomer for all the 8 clusters**. C1-8 represent clusters 1-8.

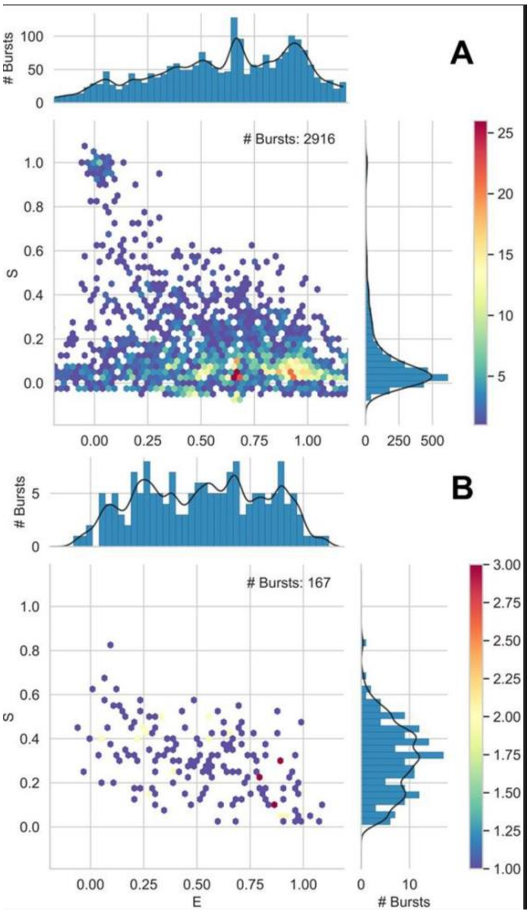


**Fig. S18: PIE-smFRET exhibit a negligible signature of dimers at picomolar concentrations.**

A mixture of donor- and acceptor-labeled αSyn (dye labeling at residue 39) at a total concentration of 100 pM, were measured for 1 hour. **A.** Selection of all bursts with a total amount of photons higher than 50 show that the majority of bursts were αSyn monomers, with apparent stoichiometry ratio (S) values close to 0 (acceptor only) and 1 (donor only). **B.** selection of all bursts that had a total of more than 25 photons arising from donor excitation and more than 25 photons arising from acceptor excitation show a few bursts with different values of mean apparent FRET efficiencies (E), and a apparent S values lower than 0.5, characteristic of dimers, before applying correction factors (after applying corrections, S should be ~0.5).

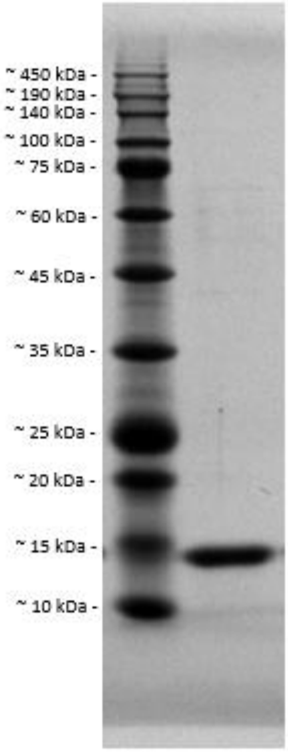


**Fig. S19: SDS-PAGE of *wt*-αSyn**

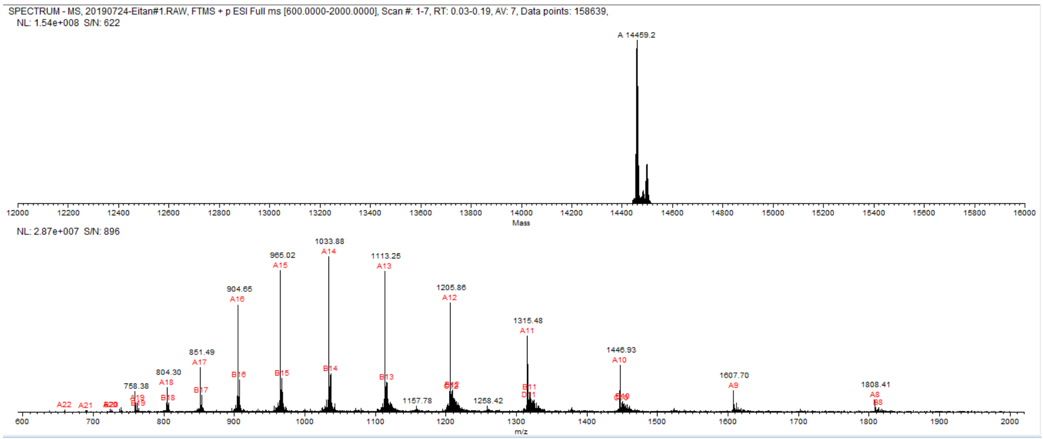


**Fig. S20: Protein intact mass determination of *wt*-αSyn**

#### Supplementary Tables:

### Table S1: Fitting sum of Gaussians to mean lifetimes histograms

| αSyn residue labeled with sCy3 | 25 pM labeled αSyn + wt αSyn Concentration (nM) | # of sub-populations | Sub-population mean lifetime (ns) | | | Fractions | | | Sub-population lifetime standard deviation (ns) | | |
|---|---|---|---|---|---|---|---|---|---|---|---|
| | | | μ1 | μ2 | μ3 | f1 | f2 | f3 | σ1 | σ2 | σ3 |
| 56 | 0 (monomer) | 3 | 2.50 ±1.04 | 4.30 ±0.29 | 6.30 ±0.67 | 0.22 ±0.29 | 0.48 ±0.56 | 0.30 ±0.63 | 1.19 ±0.43 | 1.01 ±0.16 | 1.20 ±0.14 |
| | 187.5 | 2 | 1.54 ±0.01 | 3.31 ±0.08 | | 0.66 ±0.03 | 0.34 ±0.03 | | 0.56 ±0.02 | 1.20 ±0.06 | |
| | 375 | 3 | 1.40 ±0.02 | 1.97 ±0.05 | 3.51 ±0.05 | 0.43 ±0.03 | 0.14 ±0.04 | 0.44 ±0.05 | 0.36 ±0.02 | 0.30 ±0.05 | 1.20 ±0.03 |
| | 750 | 2 | 1.58 ±0.01 | 3.46 ±0.06 | - | 0.67 ±0.02 | 0.33 ±0.02 | | 0.52 ±0.01 | 1.20 ±0.27 | |
| | 1,500 | 2 | 1.60 ±0.01 | 3.29 ±0.08 | - | 0.62 ±0.03 | 0.38 ±0.03 | | 0.53 ±0.01 | 1.20 ±0.03 | |
| 140 | 0 (monomer) | 3 | 1.67 ±0.04 | 3.50 ±0.15 | 5.33 ±0.99 | 0.20 ±0.03 | 0.70 ±0.15 | 0.10 ±0.15 | 0.65 ±0.05 | 1.20 ±0.03 | 1.20 ±0.65 |
| | 187.5 | 3 | 1.59 ±0.01 | 2.97 ±0.14 | 4.74 ±0.69 | 0.23 ±0.05 | 0.61 ±0.18 | 0.16 ±0.19 | 0.53 ±0.04 | 1.20 ±0.01 | 1.20 ±0.37 |
| | 375 | 3 | 1.49 ±0.04 | 2.84 ±0.18 | 4.63 ±0.66 | 0.14 ±0.06 | 0.65 ±0.24 | 0.22 ±0.25 | 0.51 ±0.09 | 1.20 ±0.01 | 1.20 ±0.12 |
| | 750 | 2 | 1.83 ±0.05 | 3.41 ±0.17 | - | 0.59 ±0.07 | 0.41 ±0.07 | | 0.63 ±0.05 | 1.20 ±0.08 | |
| | 1,500 | 2 | 1.78 ±0.05 | 3.53 ±0.01 | - | 0.56 ±0.05 | 0.44 ±0.05 | | 0.78 ±0.05 | 1.20 ±0.15 | |
| 39 | 0 (monomer) | 2 | 1.69 ±0.12 | 3.51 ±0.18 | - | 0.53 ±0.09 | 0.48 ±0.09 | | 0.88 ±0.06 | 1.20 ±0.11 | |
| | 187.5 | 3 | 1.72 ±0.15 | 3.29 ±0.10 | 4.73 ±0.41 | 0.28 ±0.06 | 0.39 ±0.18 | 0.33 ±0.19 | 0.88 ±0.14 | 0.84 ±0.19 | 1.20 ±0.06 |
| | 375 | 3 | 1.62 ±0.14 | 3.57 ±0.11 | 5.54 ±0.72 | 0.25 ±0.06 | 0.61 ±0.17 | 0.14 ±0.18 | 0.93 ±0.12 | 1.17 ±0.23 | 1.20 ±0.15 |
| | 750 | 3 | 1.49 ±0.08 | 3.40 ±0.08 | 5.55 ±0.66 | 0.19 ±0.03 | 0.70 ±0.11 | 0.11 ±0.11 | 0.71 ±0.09 | 1.20 ±0.16 | 1.20 ±0.06 |
| | 1,500 | 3 | 1.66 ±0.25 | 2.99 ±0.27 | 4.14 ±1.62 | 0.32 ±0.09 | 0.37 ±0.19 | 0.31 ±0.21 | 0.77 ±0.39 | 1.20 ±0.53 | 1.20 ±0.02 |
| 26 | 0 (monomer) | 3 | 1.93 ±0.13 | 3.94 ±0.06 | 6.13 ±0.51 | 0.16 ±0.04 | 0.74 ±0.11 | 0.10 ±0.12 | 0.83 ±0.14 | 1.20 ±0.21 | 1.12 ±0.72 |
| | 187.5 | 3 | 1.80 ±0.14 | 3.79 ±0.09 | 5.64 ±0.56 | 0.09 ±0.02 | 0.77 ±0.11 | 0.14 ±0.11 | 0.77 ±0.17 | 1.20 ±0.12 | 1.20 ±0.41 |
| | 375 | 3 | 1.67 ±0.08 | 3.61 ±0.20 | 5.37 ±0.76 | 0.12 ±0.03 | 0.68 ±0.21 | 0.20 ±0.21 | 0.66 ±0.10 | 1.20 ±0.63 | 1.20 ±0.16 |
| | 750 | 3 | 2.50 ±0.43 | 3.89 ±0.15 | 5.29 ±0.79 | 0.38 ±0.18 | 0.39 ±0.34 | 0.23 ± 0.38 | 1.20 ±0.07 | 0.88 ±0.25 | 1.20 ±0.13 |
| | 1,500 | 3 | 2.03 ±0.47 | 3.77 ±0.20 | 5.73 ±0.8 | 0.23 ±0.17 | 0.59 ±0.31 | 0.18 ±0.35 | 1.07 ±0.29 | 1.20 ±0.034 | 1.20 ±0.44 |

Table S2: The average pairwise RMSD (Å) for all the structures within each cluster

| | **C1** | **C2** | **C3** | **C4** | **C5** | **C6** | **C7** | **C8** |
|---|---|---|---|---|---|---|---|---|
| **RMSD (Å)** | 5.0 | 3.9 | 3.7 | 8.8 | 5.3 | 4.4 | 5.8 | 5.1 |

Table S3: The average RMSD (Å) for all the structures between each cluster

| | **C1** | **C2** | **C3** | **C4** | **C5** | **C6** | **C7** | **C8** |
|---|---|---|---|---|---|---|---|---|
| **C1** | 0 | 16.0 | 19.1 | 19.9 | 20.9 | 21.5 | 22.3 | 21.6 |
| **C2** | 16.0 | 0 | 19.2 | 20.3 | 19.3 | 19.4 | 15.9 | 20.0 |
| **C3** | 19.1 | 19.2 | 0 | 20.7 | 17.5 | 19.9 | 19.7 | 17.8 |
| **C4** | 19.9 | 20.3 | 20.7 | 0 | 20.0 | 20.3 | 19.8 | 18.6 |
| **C5** | 20.9 | 19.3 | 17.5 | 20.0 | 0 | 20.1 | 14.3 | 21.1 |
| **C6** | 21.5 | 19.4 | 19.9 | 20.3 | 20.1 | 0 | 17.3 | 20.5 |
| **C7** | 22.3 | 15.9 | 19.7 | 19.8 | 14.3 | 17.3 | 0 | 20.6 |
| **C8** | 21.6 | 20.0 | 17.8 | 18.6 | 21.1 | 20.5 | 20.6 | 0 |

Table S4: The percentage of structures with distances smaller than the cross-linking constraints for each cluster. Intermolecular cross-links 1 to 18 are $BS^3$ cross-links with a distance constraint of 30 Å. Cross-links 19 to 41 are DMTMM cross-links with a distance constraint of 16 Å.

| **Intermolecular cross-links** | **C1** | **C2** | **C3** | **C4** | **C5** | **C6** | **C7** | **C8** |
|---|---|---|---|---|---|---|---|---|
| 1: M1-K45 | 100.00 | 96.20 | 0.00 | 100.00 | 0.00 | 0.27 | 0.87 | 100.00 |
| 2: K12-K12 | 100.00 | 99.85 | 12.15 | 100.00 | 0.00 | 0.00 | 15.00 | 88.58 |
| 3: K12-K23 | 99.84 | 100.00 | 1.83 | 100.00 | 0.00 | 0.00 | 71.52 | 34.67 |
| 4: K12-K34 | 99.90 | 100.00 | 0.00 | 100.00 | 0.00 | 0.00 | 5.87 | 100.00 |
| 5: K12-K45 | 100.00 | 100.00 | 0.00 | 100.00 | 0.00 | 29.48 | 68.48 | 100.00 |
| 6: K12-K60 | 99.52 | 100.00 | 0.00 | 100.00 | 0.00 | 0.00 | 86.74 | 100.00 |
| 7: K12-K97 | 100.00 | 100.00 | 0.00 | 100.00 | 0.00 | 92.76 | 96.96 | 100.00 |
| 8: K23-K23 | 96.15 | 92.25 | 0.00 | 100.00 | 0.00 | 0.00 | 8.91 | 0.17 |
| 9: K23-K34 | 97.87 | 89.77 | 0.69 | 100.00 | 0.00 | 76.13 | 0.00 | 100.00 |
| 10: K23-K45 | 100.00 | 84.36 | 0.00 | 100.00 | 0.00 | 99.14 | 0.43 | 100.00 |
| 11: K23-K60 | 77.39 | 97.08 | 0.31 | 100.00 | 0.00 | 46.71 | 0.00 | 99.83 |
| 12: K23-K96 | 100.00 | 97.95 | 0.00 | 100.00 | 0.00 | 100.00 | 89.78 | 36.67 |
| 13: K23-K97 | 100.00 | 97.37 | 0.00 | 100.00 | 0.00 | 100.00 | 61.96 | 76.58 |
| 14: K34-K34 | 100.00 | 82.16 | 100.00 | 100.00 | 0.00 | 100.00 | 0.00 | 51.25 |
| 15: K34-K45 | 100.00 | 80.26 | 90.56 | 100.00 | 0.00 | 100.00 | 0.00 | 100.00 |
| 16: K34-K60 | 100.00 | 91.37 | 100.00 | 100.00 | 0.00 | 100.00 | 0.00 | 14.25 |
| 17: K34-K97 | 100.00 | 94.30 | 96.73 | 95.99 | 0.00 | 100.00 | 6.52 | 24.08 |
| 18: K45-K97 | 100.00 | 100.00 | 0.00 | 0.00 | 72.86 | 71.49 | 80.00 | 0.00 |
| 19: K6-E20 | 0.00 | 0.00 | 0.00 | 11.70 | 0.00 | 0.00 | 0.00 | 0.00 |
| 20: K12-E20 | 0.00 | 0.00 | 0.00 | 31.41 | 0.00 | 0.00 | 0.00 | 0.00 |
| 21: E20-K34 | 0.00 | 81.43 | 0.00 | 29.11 | 0.00 | 0.00 | 0.00 | 0.00 |
| 22: E83-K12 | 0.00 | 0.00 | 100.00 | 0.00 | 0.00 | 0.00 | 0.00 | 0.00 |
| 23: K23-E83 | 0.00 | 0.00 | 0.00 | 0.00 | 0.00 | 0.00 | 0.00 | 0.00 |
| 24: K12-E83 | 8.25 | 0.00 | 0.00 | 0.00 | 0.00 | 0.00 | 0.00 | 0.00 |
| 25: E13-K12 | 97.23 | 0.00 | 0.00 | 100.00 | 0.00 | 0.00 | 0.00 | 0.00 |
| 26: K23-E13 | 0.00 | 39.33 | 0.00 | 68.59 | 0.00 | 0.00 | 0.00 | 0.00 |
| 27: K12-E13 | 95.86 | 0.00 | 0.00 | 100.00 | 0.00 | 0.00 | 0.00 | 0.00 |
| 28: M1-E13 | 99.59 | 0.00 | 0.00 | 98.95 | 0.00 | 0.00 | 0.00 | 0.00 |
| 29: E13-K23 | 95.16 | 0.00 | 0.00 | 31.54 | 0.00 | 0.00 | 0.00 | 0.00 |
| 30: K23-E35 | 0.00 | 46.78 | 0.00 | 31.41 | 0.00 | 0.00 | 0.00 | 0.17 |
| 31: E83-K23 | 0.00 | 0.00 | 72.94 | 2.04 | 0.00 | 0.00 | 0.00 | 0.00 |
| 32: E61-K12 | 0.00 | 0.00 | 0.00 | 0.00 | 0.00 | 0.00 | 0.00 | 0.00 |
| 33: K23-E20 | 0.00 | 80.41 | 0.00 | 93.96 | 0.00 | 0.00 | 0.00 | 0.00 |
| 34: K12-E35 | 0.00 | 59.06 | 0.00 | 2.17 | 0.00 | 0.00 | 0.00 | 21.42 |
| 35: K45-E13 | 0.00 | 0.00 | 3.59 | 0.00 | 0.00 | 0.00 | 0.00 | 0.00 |
| 36: K60-E61 | 0.00 | 0.00 | 0.00 | 0.99 | 0.00 | 0.00 | 0.00 | 0.00 |
| 37: E46-K23 | 0.00 | 21.35 | 0.00 | 0.00 | 0.00 | 0.00 | 0.00 | 0.00 |
| 38: E57-K12 | 1.02 | 0.00 | 0.00 | 30.88 | 0.00 | 0.00 | 0.00 | 0.00 |
| 39: K45-E28 | 0.00 | 0.00 | 0.00 | 0.00 | 0.00 | 0.00 | 0.00 | 0.00 |
| 40: E83-M1 | 0.00 | 0.00 | 100.00 | 0.00 | 0.00 | 0.00 | 0.00 | 0.00 |
| 41: E35-K12 | 99.97 | 0.00 | 2.45 | 36.33 | 0.00 | 0.00 | 0.00 | 0.00 |

Table S5: *PPCheck*(Sukhwal & Sowdhamini, 2013)-server protein-protein interactions analyses of C1-8

| | C1 | C2 | C3 | C4 | C5 | C6 | C7 | C8 |
|---|---|---|---|---|---|---|---|---|
| H-bond[1] energy (KJ/mol) | -125.66 | -60.31 | -21.61 | -134.40 | -44.89 | -62.24 | -42.34 | -65.46 |
| Electrostatic energy (KJ/mol) | 39.57 | 8.77 | 20.62 | -31.57 | 9.19 | 0.00 | 12.66 | -0.95 |
| VdW[2] energy (KJ/mol) | -469.25 | -264.08 | -175.00 | -321.06 | -148.02 | -188.25 | -172.53 | -140.29 |
| Total stabilizing energy (KJ/mol) | -555.35 | -315.63 | -175.98 | -487.03 | -183.73 | -250.49 | -202.22 | -206.70 |
| # interface residues | 155 | 109 | 88 | 105 | 62 | 88 | 82 | 60 |
| Normalized energy per residue (KJ/mol) | -3.58 | -2.90 | -2.00 | -4.64 | -2.96 | -2.85 | -2.47 | -3.44 |
| # short contacts | 47 | 16 | 6 | 45 | 13 | 16 | 16 | 14 |
| # hydrophobic interactions | 14 | 9 | 5 | 15 | 7 | 14 | 8 | 9 |
| # VdW pairs | 19,993 | 11,955 | 7,983 | 13,956 | 5,911 | 8,405 | 8,067 | 5,550 |
| # Salt-bridges | 6 | 2 | 0 | 7 | 1 | 0 | 0 | 0 |
| # potential favorable electrostatic interactions | 14 | 8 | 5 | 12 | 2 | 0 | 1 | 3 |
| # potential unfavorable electrostatic interactions | 22 | 11 | 6 | 15 | 3 | 0 | 2 | 1 |

[1] Hydrogen-bond
[2] Van der Waals

### Table S6: β-sheet and β-strand based secondary structures

| Cluster (1-8) | Subunit (1/2) | Involving segments (NTD/NAC/CTD) | β strand #1 | β strand #2 | β strand #3 | β strand #4 |
|---|---|---|---|---|---|---|
| C1 | 1 | NTD | F4-G14 | A18-G25 | A27-K34 | - |
| C1 | 1 | NTD | V52-T54 | E57-E61 | - | - |
| C1 | 1 | NAC, CTD | G68-T81 | L100-E104 | - | - |
| C1 | 2 | NTD | V3-M5 | S9-A11 | - | - |
| C1 | 2 | NAC | A69-V71 | V77-Q79 | A85-A90 | - |
| C2 | 1 | NTD | K23-G25 | K32-K34 | - | - |
| C2 | 1 | NTD | V37-G41 | V52-A56 | - | - |
| C2 | 1 | NTD, NAC | V3-M5 | A78-V82 | G93-V95 | - |
| C2 | 1 | CTD | E131-Y133 | D135-E137 | - | - |
| C2 | 2 | NTD | K12-E20 | Q24-A30 | - | - |
| C2 | 2 | NTD | K32-K34 | L38-V40 | - | - |
| C2 | 2 | NAC | V66-G68 | V74-E83 | A85-T92 | - |
| C2 | 2 | CTD | V95-K97 | E105-A107 | - | - |
| C2 | 2 | CTD | P-117-D119 | E123-Y125 | - | - |
| C2 | 2 | CTD | P128-E130 | - | - | - |
| C2 | 2 | CTD | P138-A140 | | | - |
| C3 | 1 | NTD | M5-V15 | A17-A29 | - | - |
| C3 | 1 | NTD | G31-T33 | G36-L38 | - | - |
| C3 | 1 | NTD, NAC | V40-V55 | T59-E61 | - | - |
| C3 | 1 | NAC | G67-A69 | - | - | - |
| C3 | 1 | NAC | V77-A85 | S87-A89 | - | - |
| C3 | 1 | CTD | T92-K97 | L100-E105 | E110-I112 | - |
| C3 | 1 | CTD | P117-D119 | - | - | - |
| C3 | 1 | CTD | E123-Y125 | - | - | - |
| C3 | 1 | CTD | E130-G132 | Q134-Y136 | - | - |
| C3 | 2 | NTD | G7-S9 | - | - | - |
| C3 | 2 | NTD | K23-G25 | A27-A30 | - | - |
| C3 | 2 | NTD | K43-V48 | G51-A56 | - | - |
| C3 | 2 | NAC | K60-Q62 | - | - | - |
| C3 | 2 | NAC | T72-V74 | A76-K80 | S87-A91 | G93-V95 |
| C3 | 2 | CTD | Q103-A107 | Q109-L113 | - | - |
| C3 | 2 | CTD | A124-S129 | G132-Y136 | - | - |
| C4 | 1 | NTD | V3-M5 | - | - | - |
| C4 | 1 | NTD | K12-G14 | T22-Q24 | - | - |
| C4 | 1 | NTD | T33-E35 | Y39-G41 | K43-G47 | - |
| C4 | 1 | NTD | E57-T59 | - | - | - |
| C4 | 1 | NAC, CTD | T75-V77 | G86-I88 | G93-V95 | D98-L100 |
| C4 | 1 | CTD | E123-M127 | E131-D135 | - | - |
| C4 | 2 | NTD | E35-V40 | T44-E46 | - | - |
| C4 | 2 | NAC | A69-G73 | - | - | - |
| C4 | 2 | NAC | A78-K80 | E83-A85 | - | - |
| C4 | 2 | CTD | Q99-N103 | A107-D115 | - | - |
| C4 | 2 | CTD | Y125-S129 | G132-Y136 | P138-A140 | - |
| C5 | 1 | NTD | D2-K6 | K10-G14 | A17-A19 | - |
| C5 | 1 | NTD | K32-E35 | V37-G41 | V52-T59 | - |
| C5 | 1 | NAC | A78-V82 | S87-A90 | G93-V95 | - |
| C5 | 1 | CTD | D121-M127 | - | - | - |
| C5 | 2 | NTD | K12-G14 | V16-V19 | A27-A29 | - |
| C5 | 2 | NTD, NAC | K43-G47 | V71-G73 | A76-F94 | - |
| C5 | 2 | CTD | Q109-G111 | L113-D115 | - | - |
| C5 | 2 | CTD | E123-Y125 | - | - | - |
| C6 | 1 | NTD | V3-M5 | S9-A11 | - | - |
| C6 | 1 | NTD | G25-A27 | A30-K32 | - | - |
| C6 | 1 | NTD, NAC | K45-G47 | V49-V55 | Q79-V82 | - |
| C6 | 1 | NTD | G73-V77 | - | - | - |
| C6 | 1 | CTD | K96-D98 | K102-E104 | - | - |
| C6 | 1 | CTD | P120-Q122 | Y125-M127 | - | - |
| C6 | 2 | NTD | M5-G7 | S9-V15 | A17-K23 | A29-G31 |
| C6 | 2 | NTD | K45-T54 | - | - | - |
| C6 | 2 | NAC | V74-Q79 | - | - | - |
| C6 | 2 | NAC, CTD | S87-K96 | - | - | - |
| C6 | 2 | CTD | K102-E104 | A107-Q109 | - | - |
| C6 | 2 | CTD | M127-S129 | G132-Q134 | - | - |
| C7 | 1 | NTD | V3-K6 | K10-E13 | A17-A19 | - |
| C7 | 1 | NTD | K32-E35 | V37-G41 | V52-T59 | - |
| C7 | 1 | NAC | A78-V82 | S87-A90 | G93-V95 | - |
| C7 | 1 | CTD | E110-I112 | M116-V118 | - | - |
| C7 | 1 | CTD | D121-M127 | - | - | - |
| C7 | 2 | NTD | K10-K21 | G25-K34 | L38-V40 | - |
| C7 | 2 | NAC | V77-V82 | G86-T92 | - | - |
| C7 | 2 | CTD | G101-N103 | E105-A107 | - | - |
| C7 | 2 | CTD | P117-D119 | E123-Y125 | - | - |
| C7 | 2 | CTD | P128-E130 | - | - | - |
| C8 | 1 | NTD | M1-V3 | - | - | - |
| C8 | 1 | NTD | V15-A17 | T22-A27 | - | - |
| C8 | 1 | NTD | T33-Y39 | H50-E57 | - | - |
| C8 | 1 | NTD | K43-K45 | - | - | - |
| C8 | 1 | NAC, CTD | T64-G68 | V70-V77 | K96-D98 | - |
| C8 | 1 | NAC | K80-V82 | - | - | - |
| C8 | 1 | CTD | E123-M127 | - | - | - |
| C8 | 2 | NTD | M1-G7 | S9-E20 | T22-E28 | - |
| C8 | 2 | NTD | E35-V37 | - | - | - |
| C8 | 2 | NTD | Y39-G41 | - | - | - |
| C8 | 2 | NTD | K45-G47 | V49-V52 | - | - |
| C8 | 2 | NTD | K58-K60 | - | - | - |
| C8 | 2 | NAC, CTD | T64-V66 | A69-T72 | S87-K96 | - |
| C8 | 2 | NAC | A76-A78 | - | - | - |
| C8 | 2 | CTD | K102-E104 | A107-E110 | M116-V118 | - |
| C8 | 2 | CTD | A124-S129 | Y133-E137 | - | - |
